# Conformational Dynamics and Mechanisms of Client Protein Integration into the Hsp90 Chaperone Controlled by Allosteric Interactions of Regulatory Switches: Perturbation-Based Network Approach for Mutational Profiling of the Hsp90 Binding and Allostery

**DOI:** 10.1101/2022.05.20.492854

**Authors:** Gennady M. Verkhivker

## Abstract

Understanding allosteric mechanisms of the Hsp90 chaperone interactions with cochaperones and client protein clientele is fundamental to dissect activation and regulation of many proteins. In this work, atomistic simulations are combined with perturbation-based approaches and dynamic network modeling for a comparative mutational profiling of the Hsp90 binding and allosteric interaction networks in the three Hsp90 maturation complexes with FKBP51 and P23 cochaperones and the glucocorticoid receptor (GR) client. The conformational dynamics signatures of the Hsp90 complexes and dynamics fluctuation analysis revealed how the intrinsic plasticity of the Hsp90 dimer can be modulated by cochaperone and client protein to stabilize the closed dimer state required at the maturation stage of the ATPase cycle. In silico deep mutational scanning of the protein residues characterized the hotspots of protein stability and binding affinity in the Hsp90 complexes, showing that binding hotspots may often coincide with the regulatory centers that modulate dynamic allostery in the Hsp90 dimer. We introduce a perturbation-based network approach for mutational scanning of allosteric residue potentials and characterize allosteric switch clusters that control mechanism of cochaperone-dependent client recognition and remodeling by the Hsp90 chaperone. The results revealed a conserved network of allosteric switches in the Hsp90 complexes that allow cochaperones and GR protein become integrated into the Hsp90 system by anchoring to the conformational switch points in the functional Hsp90 regions. This study suggests that the Hsp90 binding and allostery may operate under a regulatory mechanism in which activation or repression of the Hsp90 activity can be pre-encoded in the allosterically regulated Hsp90 dimer motions. By binding directly to the conformational switch centers on the Hsp90, cochaperones and interacting proteins can efficiently modulate allosteric interactions and long-range communications required for client remodeling and activation.

## Introduction

The evolutionary conserved and conformationally adaptable Hsp90 chaperones are responsible for conformational folding of diverse client proteins and share a common homodimer topology consisting of the nucleotide-binding N-terminal domain (Hsp90-NTD), a middle domain (Hsp90-MD), involved in binding of client proteins, and a C-terminal domain (Hsp90-CTD required for constitutive dimerization.^1–8^ Structural and biophysical investigations including hydrogen/deuterium exchange mass spectrometry (HDX-MS), electron microscopy (EM) and small-angle X-ray scattering (SAXS) have detailed conformational dynamics of the Hsp90 chaperones that can spontaneously fluctuate between ensembles of open and closed conformations providing a mechanism for timely progression of the ATPase cycle and binding of client proteins.^9–15^ The functions of the Hsp90 chaperone machinery are facilitated through modulation of the conformational equilibrium by cochaperones that assist Hsp90 in orchestration of loading, processing and release of client proteins.^16,17^ The cochaperone p23 is one of the central components of the Hsp90 machinery that acts at the late stages of the ATPase functional cycle by stabilizing the ATP-bound closed form of Hsp90, inhibiting progression of the cycle, and imposing control on the time release of client proteins undergoing maturation.^18–20^ Structural studies investigated dynamics and molecular interactions of p23 with Hsp90, showing the importance of the flexible p23 tail that decelerates the ATPase by allosterically switching the conformation of the catalytic loop in Hsp90.^21^ NMR spectroscopy of the cochaperone Cdc37 interactions with Hsp90 characterized mechanisms by which this highly specialized adaptor can sense conformational instability of the kinase client proteins and load them to the chaperone system for subsequent client refolding and processing.^22–25^ NMR and HDX-MS experiments revealed how protein kinases are recruited and loaded to the Hsp90-Cdc37 complex during Hsp90-mediated chaperoning that leads to enhanced client kinase stability and activation.^25^ This study demonstrated a critical importance of conformational dynamics for the client loading and remodeling mechanism that is facilitated by dynamic sampling of low-populated conformational states of all interacting partners and enabled by a cascade of allosteric conformational switches at the binding interfaces. The pioneering cryo-electron microscopy structure of the Hsp90-Cdc37-Cdk4 kinase complex has provided the full atomistic details of the Hsp90 interactions with cochaperone and Cdk4 client protein, revealing a complex and dynamically-controlled mechanism that involves allosteric cooperativity and remodeling of spatially separated binding interfaces.^26,27^ Another important cochaperone Aha1 accelerates the ATPase cycle by biasing Hsp90 towards an N-terminally closed ATP-bound conformation immediately preceding ATP hydrolysis.^28^ NMR studies showed that a single Hsp90-MD residue Y313 is a phosphorylation-sensitive conformational switch that affects both long-range conformational dynamics to facilitate the recruitment of Aha1 to the Hsp90 system.^29^ Biophysical and NMR studies of the Hsp90 complexes further characterized the diversity and plasticity of multiple binding interfaces available for protein clients on Hsp90 by examining chaperone interactions with the ligand-binding domain (LBD) of the glucocorticoid receptor (GR),^30^ the DNA binding domain (DBD) of p53 client,^31^ and the intrinsically disordered Tau protein.^32–34^ NMR spectroscopy, and chemical cross-linking analysis of the Hsp90-FKBP51 complex with Tau protein showed evidence of dynamic allostery, in which FKBP51 stabilizes the open conformation of the Hsp90 to facilitate Tau binding and the inherent flexibility of the disordered Tau protein is precisely coupled to the dynamic changes in the Hsp90 chaperone^34^. Subsequent biochemical and NMR studies established that FKBP51 cochaperone protein can cooperate with p23 during client maturation.^35,36^

The cryo-EM structure of the human Hsp90-FKBP51-p23 complex disclosed a complex molecular basis of cochaperone function during client maturation, in which FKBP51 cochaperone recognizes a closed Hsp90 conformation via binding to Hsp90-MD/CTD near the client binding sites, whereas a single p23 molecule allosterically stabilizes the closed dimer (Figure 1A,B).^37^ In the FKBP51 protein, the N-terminal FK1 domain is connected to a related but inactive FK2 domain followed by the C-terminal tetratricopeptide-repeats (TPR) domain, which together adopt an extended tripartite configuration.^38,39^ In the Hsp90-FKBP51-P23 complex, the TPR C-terminal extension helix (H7e) of FKBP51 recognizes the Hsp90-CTD functional regions using a conformation different from the crystal structures,^38,39^ while the FK1 domain is positioned near Hsp90-MD client binding site to enable cochaperone-induced modulation of the Hsp90-client interactions (Figure 1A,B). The biochemical analysis of the Hsp90-FKBP51-P23 structure also suggested a mechanism in which the TPR binding interactions of FKBP51 with the Hsp90-CTD can be allosterically coupled with a more dynamic FK1-Hsp90/MD binding interface and together with P23 cochaperone that enforces the closed Hsp90 conformation allow to accelerate GR client remodeling on the Hsp90 (Figure 1A,B).^37^ The complexity and diversity of the Hsp90-client interactions was recently illuminated by Agard and colleagues in the cryo-EM structure of the Hsp90-Hsp70-Hop-GR-loading complex that disclosed an unexpected mode of coordination between the Hsp90 and two Hsp70 proteins, in which one Hsp70 delivers GR client protein to the chaperone system, while the second Hsp70 molecule serves as scaffolding platform for the Hop cochaperone.^40^ The accompanied study of the maturation Hsp90-p23-GR complex presented the first atomic resolution structure of an activated and folded GR client bound to the Hsp90, in which p23 cochaperone stabilizes the closed Hsp90 dimer and interacts with the GR client protein.^41^

**Figure 1.**
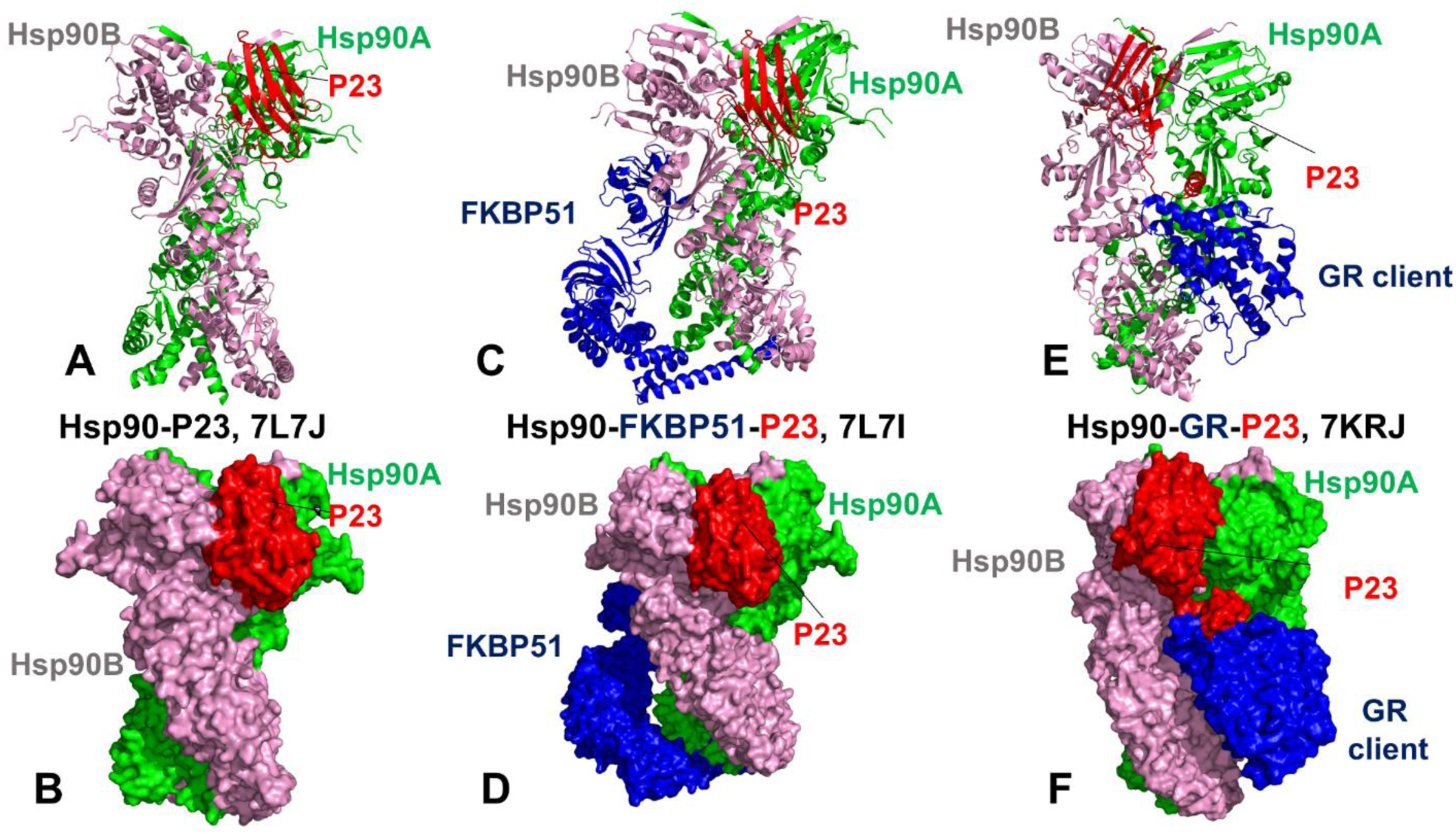
Structural Organization of the Hsp90 complexes. The structure of the human Hsp90-P23 complex is shown in ribbons (A) and surface representation (B). The structure of the human Hsp90-FKBP51-P23 complex (C,D). The structure of the human Hsp90-GR-P23 complex (E,F). The Hsp90-A monomer of the dimer is shown in green, Hsp90-B is shown in pink, P23 is depicted in red, FKBP51 cochaperone and GR client protein are shown in blue.

These pioneering studies have opened up a new chapter in the chaperone research by mapping out at the atomic resolution the chaperone-mediated client loading and activation mechanisms of the complete Hsp90 chaperone cycle, and revealing how structural adaptability of the binding partners can be exploited to ensure a protected dynamic refolding of the Hsp90 clients and subsequent stabilization of the activated folded client by specific interactions with P23 cochaperone.^41^ These studies highlighted the evolving diversity and dynamics of the Hsp90-cochaperone-client arrangements in which the allosteric cooperativity between distinct binding interfaces formed by Hsp90 and cochaperones that can be regulated through spatially separated conformational switch centers. NMR spectroscopy and SAXS experiments examined the Hsp90 binding with globular and partially disordered clients, showing that client binding can induce allosteric population shifts between the Hsp90 states as well as dynamic allosteric changes in the closed Hsp90 dimer that are propagated across chaperone domains through conserved allosteric network of communication switches.^42^ Single molecule Förster resonance energy transfer (smFRET) experiments disentangled important drivers underlying allosteric modulation of the Hsp90 conformational dynamics that allow to constrain the catalytically active elements and promote the preferential stabilization the closed dimer through several mechanisms including site-specific mutations of conformational switch centers, via cochaperone binding, and through non-specific macro-molecular crowding.^43^ Nanosecond single-molecule fluorescence fluctuation analysis of the Hsp90 dynamics combined with mutagenesis experiments detected allosteric communication between structural rearrangements at remote positions of the Hsp90 dimer that remained mobile on a sub-millisecond time scale.^44^ The domain motions within the apo-state of the Hsp90, probed by gold nano-spheres revealed long-range allosteric cooperativity that occurs on a broad range of time scales ranging from seconds to minutes^45^ suggesting that the intrinsic dynamic landscape of the Hsp90 dimer can enable a diverse range of fine-tuning and functional modulation responses by cochaperones and client proteins. The advanced fluorescence methods combined with molecular dynamics (MD) simulations examined ATP-induced allosteric changes in the Hsp90 that involve hierarchical dynamics on timescales from nano-to milliseconds.^46^ Two-color fluorescence microscopy in combination with photo-induced electron transfer (PET) also observed that the Hsp90 structural changes can occur cooperatively, revealing synchronicity and allosteric coupling of conformational motions at remote sites during ATPase-

driven closure of the Hsp90 dimer in the chaperone’s catalytic cycle.^47^ In summary, a significant body of biophysical and structural studies have demonstrated that the Hsp90 chaperone can function as allosterically regulated adaptative machine that fluctuates between distinct functional forms, where the interactions with cochaperones and client proteins can allosterically modulate population shifts and dynamic changes in the functional states required for timely processing of folded clients. A number of single conformational switches in the Hsp90 can control population shifts between open and closed Hsp90 states, modulation through cochaperone recruitment and client binding. The Hsp90 interactions can be regulated by phosphorylation sites^48–52^ and their functional role as allosteric switches of allosteric conformational changes during the ATPase cycle was demonstrated.^53–57^ The recruitment of Aha1 co-chaperone to Hsp90 can be enhanced by phosphorylation of Y313 in Hsp90α (Y305 in Hsp90β, Y293 in Hsp82) in the Hsp90-MD region.^29,58^ A multi-purpose molecular switch W300 in the Hsp82-MD (W320 in human Hsp90α) is required for chaperone regulation and allows to exert control over the conformational changes and timely access to the client-competent conformations during the ATPase cycle.^59^ Mutations of the W320 conformational switch result in significant deleterious effects impeding GR client proper processing and activation.^60^ This regulatory center is also instrumental for integration of the Aha1 cochaperone to the chaperone system^61^ highlighting multi-purpose role of W320 for anchoring both clients and cochaperones. A conserved methylated lysine site in the CTD K615 of Hsp90α (K607 for Hsp90β and K594 for Hsp82) is an important switch point that allosterically regulates the ATPase activity, dimer closing kinetics, co-chaperone modulation, and client maturation of the Hsp90.^62^

Computational studies and atomistic MD simulations of the Hsp90 chaperones have provided evidence for allosteric communications in the Hsp90 dimer^63–67^ and enabled computer-based discovery of allosteric modulators of the chaperone functions.^68–73^ Integration of molecular simulations and network-based modeling of the Hsp90 interactions with cochaperones and client proteins suggested that allosteric coupling between key functional centers may be utilized in regulation of the inter-domain communications, control of ATP hydrolysis, and protein client binding.^74–81^ Using molecular simulations and perturbation-based network modeling of the Hsp90-Cdc37 complexes with kinase clients, we characterized the mechanism of phosphorylation-induced communication switching.^82^ By simulating the effect of cochaperones and client proteins on signal transmission in the Hsp90 chaperone system, we suggested that Hsp90 interactions may be controlled by the clusters of regulatory control points that are assembled in structural communication spines that mediate long-range interactions in the Hsp90 chaperone.^83^ An allosteric mechanism of asymmetric ATP binding in the Hsp90 and its implications for client remodeling was recently reported showing the existence of regulatory control centers acting over the long-range to modulate chaperone function.^84^ In our latest study, we simulated the cryo-EM structure of the Hsp90-Hsp70-Hop-CR client complex and employed perturbation-based network modeling methods to show that the Hsp90 interactions can be governed by two allosteric residue clusters that exploit the intrinsically wired Hsp70 allostery to mediate integration of the Hsp70-bound client into the Hsp90 chaperone system.^85^

In the current work, we expand on that study by performing a comparative computational analysis of the Hsp90-P23, Hsp90-FKBP51-P23 and Hsp90-GR-P23 complexes (Figure 1) to investigate dynamic chaperone interactions and identify key regulatory centers that control mechanism of cochaperone-dependent client recognition and remodeling. The ensemble-based distance fluctuation analysis quantifies the effect of cochaperone P23 and GR client on the Hsp90 dynamics, showing that protein binding can curtail coordinated movements of the Hsp90 dimer and slow down the inter-domain allosteric signals, which consistent with the functional role of P23 in arresting the ATPase cycle and stabilizing the closed dimer state. Using systematic mutational scanning of protein residues and mutational cartography analysis, we characterize the hotspots of protein stability and binding affinity in the Hsp90 complexes, particularly showing that W320 switch is also the most important binding affinity hotspot critical for processing of the GR client protein. Allosteric communication between distant sites in proteins is central to biological regulation but still poorly characterized due to limitations of methods to comprehensively quantify allostery in diverse proteins. Here we attempt address some shortcomings by presenting an in silico analog of “deep” mutational scanning to globally map allosteric preferences.

We introduce perturbation-based network approach for mutational scanning of allosteric residue preferences and determine allosteric hotspots whose mutations result in dramatic impairment of the long-range communications in the interaction network. The results reveal conformational switches of the Hsp90 dynamics that support a mechanism, according to which P23 and GR proteins are integrated into Hsp90 system by anchoring to the conformational switch points in the MD and CTD regions. By directly interacting with conformational CTD switch region, GR binding can impose dynamic constraints on allosteric communication between Hsp90 domains and together with P23 stabilize the conformational Hsp90 form required for the maturation of client proteins. This study suggests that the Hsp90 interactions with client proteins may operate under dynamic-based allostery in which activation or repression of the Hsp90 activity can be pre-encoded in the Hsp90 dimer and preexisting conformational switches are activated through client-specific interactions.

## Materials and Methods

### Structure Analysis

The Hsp90-P23, Hsp90-FKBP51-P23 and Hsp90-GR-P23 structures were obtained from the Protein Data Bank.^86^ During structure preparation stage, protein residues in the crystal structures were inspected for missing residues and protons. Hydrogen atoms and missing residues were initially added and assigned according to the WHATIF program web interface.^87,88^ The structures were further pre-processed through the Protein Preparation Wizard (Schrödinger, LLC, New York, NY) and included the check of bond order, assignment and adjustment of ionization states, formation of disulphide bonds, removal of crystallographic water molecules and co-factors, capping of the termini, assignment of partial charges, and addition of possible missing atoms and side chains that were not assigned in the initial processing with the WHATIF program. The missing loops in the studied cryo-EM structures of the Hsp90-Hsp70-Hop-GR complex were reconstructed and optimized using template-based loop prediction approaches ModLoop,^89^ ArchPRED Server^90^ and further confirmed by FALC (Fragment Assembly and Loop Closure) program.^91^ The side chain rotamers were refined and optimized by SCWRL4 tool.^92^ The protein structures were then optimized using atomic-level energy minimization using 3Drefine method.^93^

### Coarse-Grained Simulation Models

Conformational dynamics of the Hsp90 complexes was explored using high-resolution, coarse-grained (CG) simulations within the CABS-flex approach followed by MODELLER-based full atomistic reconstruction of the resulting trajectories.^94–99^ This approach employs a high-resolution coarse-grained model and an efficient Monte Carlo conformational search which allow for simulations of large biomolecules on long time scales. In the CG-CABS model, the amino acid residues are represented by Cα, Cβ, the center of mass of side chains and another pseudoatom placed in the center of the Cα-Cα pseudo-bond. The sampling scheme of the CABS model is based on Monte Carlo replica-exchange dynamics and involves a sequence of local moves of individual amino acids in the protein structure as well as moves of small fragments.^94,95^ CABS-flex standalone dynamics package implemented in Python 2.7 was used for fast simulations of protein structures.^96–99^ The default settings were applied in which soft native-like restraints are imposed only on pairs of residues fulfilling the following conditions : the distance between their *C*^α^ atoms was smaller than 8 Å, and both residues belong to the same secondary structure elements. A total of 100 independent CG-CABS simulations were performed for the Hsp90 complexes. In each simulation, the total number of cycles was set to 1,000 and the number of cycles between trajectory frames was 100. The all-atom reconstruction CG-CABS trajectories was done using MODELLER-based reconstruction of simulation trajectories.

### MD Refinement Simulations of the Hsp90 Complexes

Multiple CG-CABS simulations of the Hsp90 complexes are followed by all-atom MD-based refinement in which we leveraged the use of multiple shorter trajectories to facilitate the broader exploration of conformational space and enable a sufficiently adequate representation of the conformational landscapes. The MD stage consisted of taking the average models from twenty different CG-CABS simulations followed by 50 ns of all-atom MD refinement simulations in explicit solvent. MD simulations were performed for an N, P, T ensemble in explicit solvent using NAMD 2.13 package^100^ with CHARMM36 force field.^101^ Long-range non-bonded van der Waals interactions were computed using an atom-based cutoff of 12 Å with switching van der Waals potential beginning at 10 Å. Long-range electrostatic interactions were calculated using the particle mesh Ewald method^102^ with a real space cut-off of 1.0 nm and a fourth order (cubic) interpolation. Energy minimization after addition of solvent and ions was conducted using the steepest descent method for 100,000 steps. All atoms of the complex were first restrained at their crystal structure positions with a force constant of 10 Kcal mol^-1^ Å^-2^. Equilibration was done in steps by gradually increasing the system temperature in steps of 20K starting from 10K until 298 K and at each step 1ns equilibration was done keeping a restraint of 10 Kcal mol-1 Å-2 on the protein C_α_ atoms. After the restrains on the protein atoms were removed, the system was equilibrated for additional 10 ns. An NPT production simulation was run on the equilibrated structures for 50 ns keeping the temperature at 310 K and constant pressure (1 atm). In simulations, the Nose–Hoover thermostat^103^ and isotropic Martyna–Tobias–Klein barostat^104^ were used to maintain the temperature at 298 K and pressure at 1 atm respectively. Principal component analysis (PCA) of MD trajectories was conducted based on the set of backbone heavy atoms using the CARMA package.^105^ The relative solvent accessibilities (RSA) of protein residues were obtained using web server GetArea.^106^

### Distance Fluctuations Stability Analysis

We employed distance fluctuation analysis of the simulation trajectories to compute residue-based stability profiles. The fluctuations of the mean distance between each pseudo-atom belonging to a given amino acid and the pseudo-atoms belonging to the remaining protein residues were computed. The fluctuations of the mean distance between a given residue and all other residues in the ensemble were converted into distance fluctuation stability indexes that measure the energy cost of the residue deformation during simulations.^107–110^ The distance fluctuation stability index for each residue is calculated by averaging the distances between the residues over the simulation trajectory using the following expression:

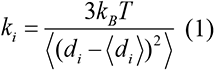

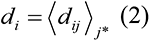

*^d^_ij_* is the instantaneous distance between residue *I* and residue *j*, *k_B_* is the Boltzmann constant, *T* =300K. denotes an average taken over the MD simulation trajectory and 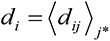 is the average distance from residue *I* to all other atoms *j* in the protein (the sum over *j*_*_ implies the exclusion of the atoms that belong to the residue *i*). The interactions between the *C_α_* atom of residue *i* and the *C_α_* atom of the neighboring residues *i* -1 and *i* +1 are excluded in the calculation since the corresponding distances are constant. The inverse of these fluctuations yields an effective force constant *k_i_* that describes the ease of moving an atom with respect to the protein structure. The dynamically correlated residues whose effective distances fluctuate with low or moderate intensity are expected to communicate over long distances with the higher efficiency than the residues that experience large fluctuations. Our previous studies showed that residues with high value of these indexes often serve as structurally stable centers and regulators of allosteric signals, whereas small values of the distance fluctuation stability index are typically indicative of highly dynamic fluctuating sites.^77^ The structurally stable and densely interconnected residues as well as moderately flexible residues that serve as a source or sink of allosteric signals could feature high value of these indexes.

### Protein Stability and Binding Free Energy Calculations

We employed two complementary approaches to compute protein stability changes and perform mutational sensitivity analysis using systematic scanning of the intermolecular interface residues for the ensembles of the Hsp90 complexes. The FoldX approach with the all-atom representation of protein structure was used to conduct alanine scanning of the interfacial residues and evaluate protein stability changes.^111–115^ The protein stability ΔΔG changes were computed by averaging the results of computations over 1,000 samples obtained from MD simulation trajectories.

In addition, the protein-protein interfaces in the Hsp90-Hsp70-Hop-GR conformational ensemble were systematically modified and the resulting binding free energy changes were evaluated coarse-grained BeAtMuSiC approach that utilizes statistical potentials that describe pairwise inter-residue distances, backbone torsion angles and solvent accessibilities derived from known protein structures.^116–118^. BeAtMuSiC approach considers the effect of the mutation on both the strength of the interactions at the interface and on the overall stability of the complex. The binding free energy of protein-protein complex can be expressed as the difference in the folding free energy of the complex and folding free energies of the two protein binding partners:

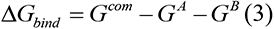

The change of the binding energy due to a mutation was calculated then as the following:

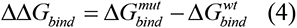

### Dynamic-Based Modeling of Residue Interaction Network and Community Analysis

A graph-based representation of protein structures^119,120^ is used to represent residues as network nodes and the inter-residue edges to describe non-covalent residue interactions. The network edges that define residue connectivity are based on non-covalent interactions between residue side-chains that define the interaction strength *I_ij_* as used in the original studies.^119,120^ The residue interaction networks are constructed using both dynamic correlations^121^ and coevolutionary residue couplings^122^ that yield robust network signatures of long-range couplings and communications. The details of this model were described in our previous studies.^123,124^ In our model, the dynamics of the system is reduced to a network of correlated local motions, and signal propagation is described as an information exchange through the network.

The RING program^125,126^ was also employed for the initial generation of residue interaction networks. The ensemble of shortest paths is determined from matrix of communication distances by the Floyd-Warshall algorithm.^127^ Network graph calculations were performed using the python package NetworkX.^128^ The Girvan-Newman algorithm^129–131^ is used to identify local communities. In this approach, edge centrality (also termed as edge betweenness) is defined as the ratio of all the shortest paths passing through a particular edge to the total number of shortest paths in the network. The method employs an iterative elimination of edges with the highest number of the shortest paths that go through them. An improvement of Girvan-Newman method was implemented, and the algorithmic details of this modified scheme were given in our recent studies.^132,133^ The betweenness of residue *i* is defined as the sum of the fraction of shortest paths between all pairs of residues that pass through residue *i* :

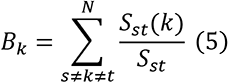

where *S_st_* denotes the number of shortest geodesics paths connecting s and *t,* and *S_st_*(*k*) is the number of shortest paths between residues *s* and *t* passing through the node *k*.

The following Z-score is then calculated as

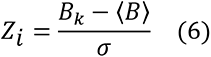

Through mutation-based perturbations of protein residues we compute dynamic couplings of residues and changes in the average short path length (ASPL) averaged over all modifications in a given position. The change of ASPL upon mutational changes of each node is inspired and reminiscent to the calculation proposed to evaluate residue centralities by systematically removing nodes from the network.

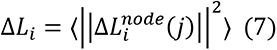

where *i* is a given site, *j* is a mutation and 〈⋯〉 denotes averaging over mutations. 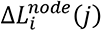 describes the change of ASPL upon mutation *j* in a residue node *i*. Δ*L_i_* is the average change of ASPL triggered by complete mutational profiling of this position.

Z-score is then calculated for each node as follows:

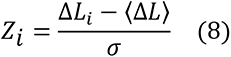

〈Δ*L*〉 is the change of the ASPL under mutational scanning averaged over all protein residues in the S-RBD and σ is the corresponding standard deviation. The ensemble-averaged Z –scores ASPL changes are computed from network analysis of the conformational ensembles using 1,000 snapshots of the simulation trajectory for the native protein system.

## Results and Discussion

### Structural Analysis and Topography of the Hsp90 Regulatory Complexes

The cryo-EM structures of the human Hsp90-FKBP51-P23 and Hsp90-GR-P23 complexes highlighted a similar topology of the Hsp90 closed dimer and diverse arrangement of the Hsp90 intermolecular interfaces (Figure 1). The conformations of the closed Hsp90 dimer in the Hsp90-P23 complex and Hsp90-FKBP51-P23 complexes (Figure 2) are similar to the ATP-bound structures of yeast Hsp90 complex with P23 cochapeone^134^ ( the root-mean-square deviation, RMSD=1.723 Å) and human Hsp90 dimer in the Hsp90-Cdc37-Cdk4 complex^26^ (RMSD=1.284 Å). In the Hsp90-P23 and Hsp90-FKBP1-P23 complexes the conserved F103 and W106 of p23 pack into the hydrophobic pocket of Hsp90-MD (Figure 2). These residues and interactions are highly conserved in yeast Hsp82-P23-Sba1 complex.^134^ The FKBP51 helix H7e residues Y409, M412 an F413 that are conserved among the immunophilin class of TPR cochaperones form a conserved hydrophobic interface with the Hsp9-CTD helices (Figure 2) which is believed to represent a conserved Hsp90 recognition mechanism for these cochaperones.^37^ In the Hsp90-GR-P23 complex, P23 cochaperone similarly interacts with the Hsp90-B protomer using conserved residues F103, N104 and W106 (Figure 3). The evolutionary and structural conservation of this Hsp90-P23 interface suggested that these interactions could play an important role in stability, binding and regulation of P23-mediated chaperone functions. The Hsp90-GR-P23 complex revealed several major interfaces formed by Hsp90 and GR client protein, including (a) Hsp90B interactions with the pre-helix 1 region of GR client (residues 523-531); (b) the interface between GR helix 1 ( residues 532–539) packing up against the amphipathic helical hairpin of Hsp90 B (residues 607–621); and (c) Hsp90A-MD binding interface residues with GR client using the conserved hydrophobic residues F349 and W320 (Figure 3).

**Figure 2.**
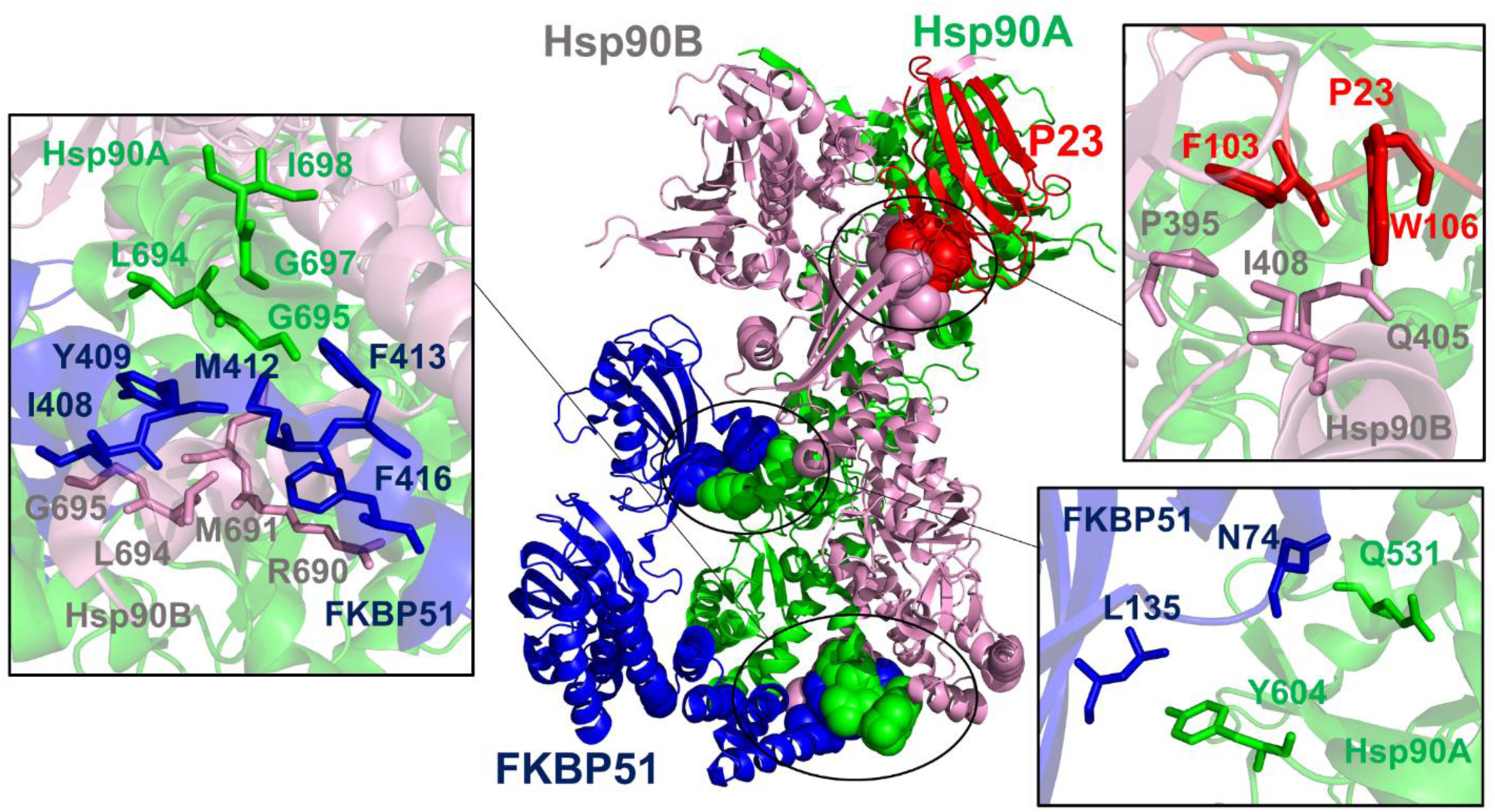
Structural analysis of the Hsp90-FKBP51-P23 complex and intermolecular interfaces. (A) A general structural overview and mapping of the intermolecular interfacial residues in the Hsp90-FKBP51-P23 regulatory complex. The Hsp90-A is in green, Hsp90-B is in pink, P23 cochaperone is in red, and FKBP51 cochaperone is in blue ribbons. The interfacial residues are shown in spheres colored according to the respective protein component. The main clusters of the intermolecular interactions are annotated by circles and pointed by arrows. (B) The primary FKBP51 recognition interface in which conserved residues Y409, M412, F413, F416 of FKBP51 interacting with the Hsp90-CTD helices (residues 658-696). FKBP51 residues are in blue sticks, Hsp90A-CTD residues are in green sticks and Hsp90B-CTD residues are in pink sticks. (C) The primary P23 binding interface with Hsp90B-MD. The conserved F103 and W106 of p23 (in red sticks) bind Hsp90-MD residues P395, Q405, I408, and V411 (in pink sticks). (C) The secondary FKBP51 binding interface (D72, N74, L135 in blue sticks) interacting with Hsp90A-MD client binding site (residues Q531, Y604 in green sticks).

**Figure 3.**
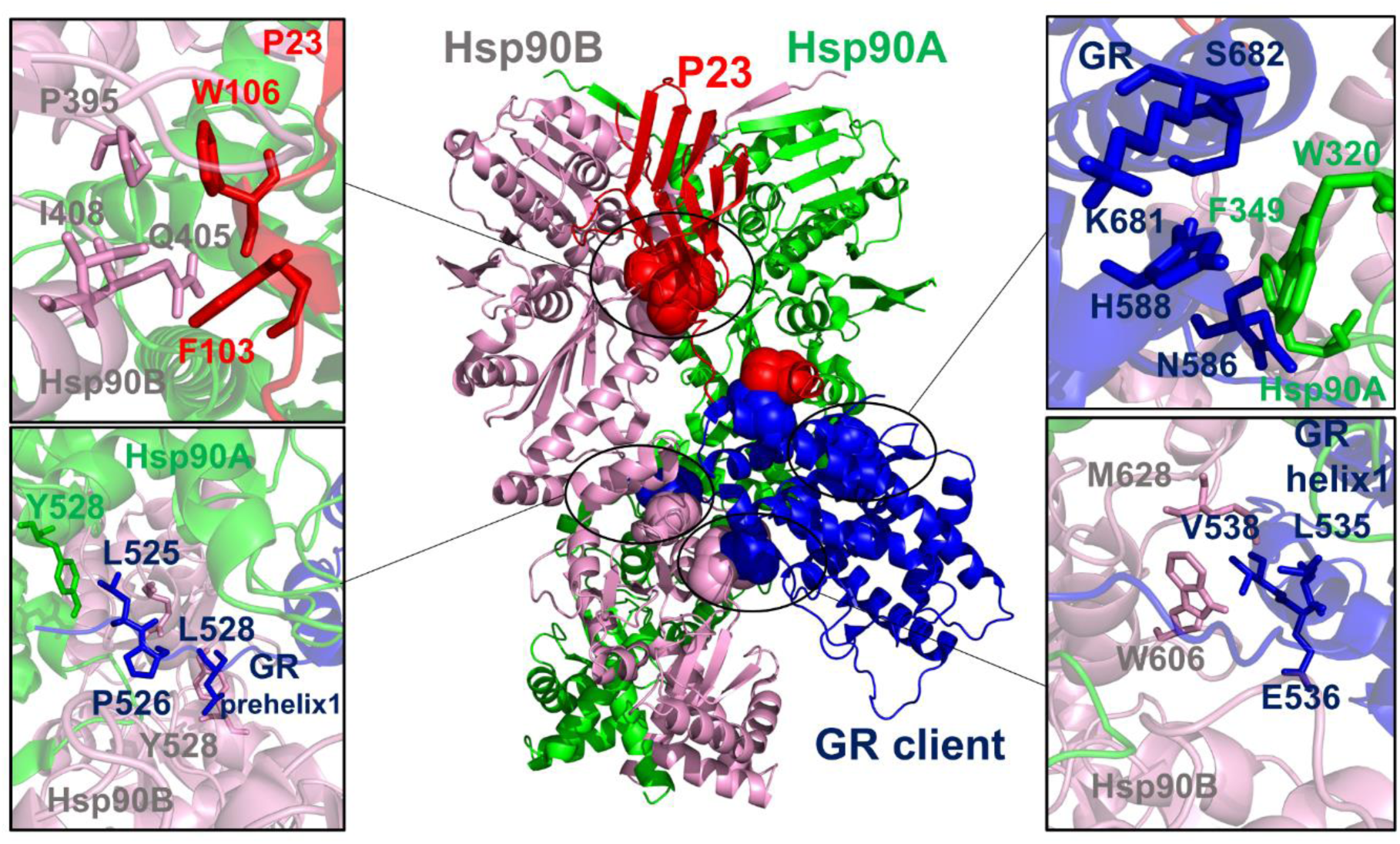
Structural analysis of the Hsp90-GR-P23 complex and intermolecular interfaces. (A) A general structural overview and mapping of the intermolecular interfacial residues in the maturation Hsp90-GR-P23 regulatory complex. The Hsp90-A is in green, Hsp90-B is in pink, P23 cochaperone is in red, and GR client protein is in blue ribbons. The interfacial residues are shown in spheres colored according to the respective protein component. The main clusters of the intermolecular interactions are annotated by circles. (B) The conserved P23 binding interface with Hsp90B-MD. The conserved F103 and W106 of p23 (in red sticks) bind Hsp90-MD residues P395, Q405, I408, and V411 (in pink sticks). (C) The binding interface between GR pre-helix 1 region (residues L525, P526, L528 in blue sticks) and Hsp90 internal lumen site (residues Y528 and Q531 in green and pink sticks for Hsp90-A and Hsp90-B respectively). (D) The binding interface between conformational switches on the Hsp90-A (W320, F349 in green sticks) and GR client (residues N586, H588, K681, S682 in blue sticks). (E) The intermolecular interface between Hsp90B-CTD amphipathic helical hairpin region (Y604, W606, M628 in pink sticks ) and GR client helix 1 ( residues L535, E536, and V538 in blue sticks).

The survey of the Hsp90 interactions with P23, FKBP51 and GR proteins indicated that the interacting proteins could exploit conserved recognition elements and conformational switches along with diversity and modular nature of the binding interfaces to enable cochaperone-mediated binding, processing and release of protein clients.

### Conformational Dynamics of the Hsp90 Regulatory Complexes: Balancing Protein Stability and Conformational Adaptability for Client Recruitment and Loading

We employ a range of computational approaches to examine the molecular details underlying conformational dynamics, binding and regulation of the Hsp90 complexes, assuming that dynamic allostery and cooperativity of the intermolecular interfaces may be an important underlying mechanism of the Hsp90 functional cycle. Conformational dynamics of the Hsp90 complexes was explored using high-resolution, coarse-grained (CG) simulations within the CABS-flex approach followed by MODELLER-based full atomistic reconstruction of the resulting trajectories. Multiple CG-CABS simulations of the Hsp90 complexes are followed by all-atom MD-based refinement in which we leveraged the use of multiple shorter trajectories to facilitate the broader exploration of conformational space and enable a sufficiently adequate representation of the conformational landscapes. The equilibrium conformational ensembles were examined using distance fluctuations coupling analysis which is based on the dynamic residue correlations ^107–110^ (Figure 4). In this model, the fluctuations of the mean distance between a given residue and all other residues in the ensemble are converted into distance fluctuation stability indexes where residues with high value of these indexes correspond to structurally stable sites. We focused on changes in the conformational variations of the Hsp90 chaperone and the effect of binding on changes in the dynamic fluctuation. First, we compared the distance fluctuation profiles for the Hsp90 protomers in the Hsp90-P23 complex (Figure 4A), Hsp90-FKBP51-P23 complex (Figure 4B) and Hsp90-GR-P23 assembly (Figure 4C). The overall shape of the profiles is determined by the conserved topology of the closed Hsp90 dimer and is similar in all complexes. The highest peaks are broadly concentrated in the Hsp90-MD regions (residues 330-450) featuring a considerable number of structurally stable sites (Figure 2A-C). The central three-helix Hsp90-MD bundle (residues 406-462, helix 1: residues 406-428; helix 2: residues 430-451; helix3: residues 455-462) links the inter-domain regions and harbors most of the high stability Hsp90 residues. In particular, the high values of distance fluctuation stability parameter were observed for Hsp90-MD residues 355-360 and residues 441-445 from helix 3 of the 3-helix bundle (Figure 4A-C). At the same time, the helix 2 residues 406-428 displayed smaller fluctuation stability indexes and are more dynamic. This helix packs across the entire central β-sheet of the Hsp90-MD, and thus can affect the dynamics of the Hsp90 near NTD-MD interdomain regions. It is worth noting that fluorescence methods identified this segment as an important allosteric mediator region that can modulate dynamics of the client binding site.^46^ A series of smaller stability peaks was seen in the Hsp90-NTD dimerization interface is composed of the first helix (residues 23-36) and the helical lid motif (residues 105-139). These motifs are known as molecular switches of Hsp90 dimerization and corresponded to the distribution peaks in all Hsp90 complexes (Figure 4A-C). The Hsp90-MD amphipathic loop (residues 349–359) and Hsp90-CTD amphipathic helical hairpin (residues 607–621) are the key functional regions that mediate dimerization and allosteric communications between NTD and CTD regions.^13,14^

**Figure 4.**
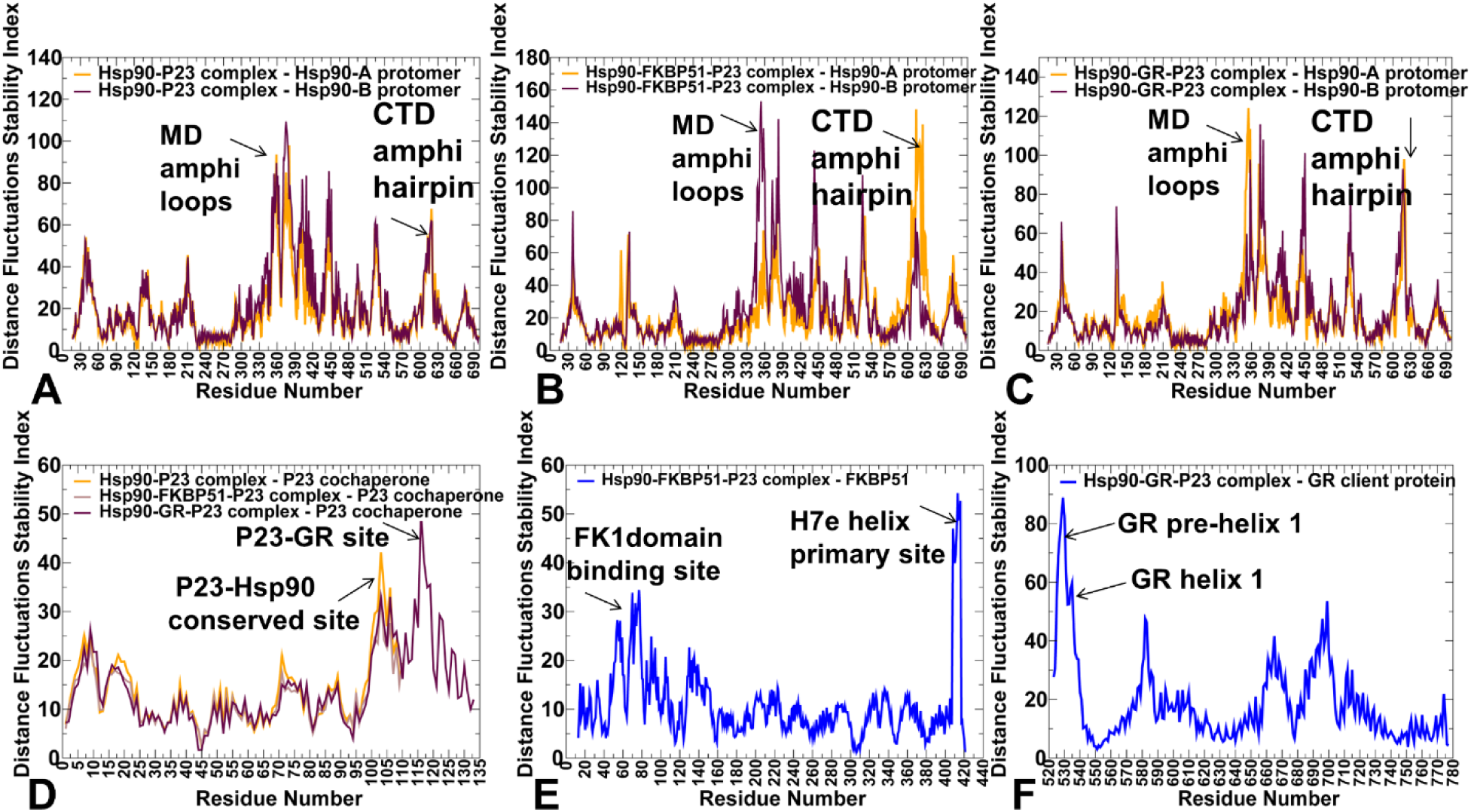
Conformational dynamics of the Hsp90 complexes. The residue-based distance fluctuation stability index for the Hsp90 dimer protomers in the Hsp90-P23 complex (A), Hsp90-FKBP51-P23 complex (B), and Hsp90-GR-P23 complex (C). The profile for the Hsp90-A protomer is in orange lines and Hsp90-B protomer in maroon-colored lines. The distance fluctuation stability index for P23 cochaperone residues in the Hsp90-P23 ( in orange), Hsp90-FKBP51-P23 (in light brown), and Hsp90-GR-P23 (in maroon) (D). The distribution for FKBP51 residues (in blue) in the Hsp90-FKBP51-P23 (E), and the profile for GR client protein ( in blue) in the Hsp90-GR-P23 complex. The important peaks of the distance fluctuation stability index are annotated on panels (A-F) and pointed by arrows.

The consistent peaks of the distribution were aligned with the regulatory Hsp90-CTD residues 607-621 (Figure 4A-C). Moreover, the relative contribution of this region and corresponding stability values increased in the Hsp90-FKBP51-P23 (Figure 4B) and Hsp90-GR-P23 complexes (Figure 4C), indicating that FKBP51 cochaperone and GR client binding may induce stabilization of these amphipathic helices. Notably, the Hsp90-CTD amphipathic helix region has been previously shown to binding client proteins in the closed state^135^ and is also utilized for Aha1 cochaperone binding.^61^ In addition, notable peaks were observed for two CTD helices (αH17 and αH18, residues 657–695) that are known to be essential for Hsp90 dimerization and hydrolysis and correspond to regulatory switch regions responsible for modulation of Hsp90 conformational changes.^13,14^ Importantly, single-molecule experiments established that the ATP-modulated dynamic opening and closing of the Hsp90-CTDs are anticorrelated with the movements of the Hsp90-NTDs, and allosteric communication between the Hsp90 domains is regulated by dynamics of the CTD αH17 and αH18 helices.^13^ The emergence of these CTD regions as critical clusters of structural stability in the Hsp90 complexes is consistent with differential scanning fluorimetry and multi-angle light scattering experiments showing that mutations in these CTD helices caused a significant decrease in the melting temperature indicating a reduced stability of the mutant Hsp90 complexes due to the disruption of the dimerization interface.^136^ The Hsp90-p23 contacts are conserved between yeast and human Hsp90 systems. P23 binding stabilizes Hsp90-NTD interacting residues I34, K41, M119, E120, A124, and A126 residues that form a noticeable peak in all distributions (Figure 4D). p23 binding imposed constrains on the mobility of the Hsp90 mobility, which is particularly evident in the Hsp90-FKBP51-P23 complex where FKBP51 and P23 rigidify the Hsp90 dimer in the MD (residues 282-526) and CTD regions (residues 547-698) (Figure 4B). In particular, the interactions of P23 with D393, O395, Q405, I408, and V411 residues of the Hsp90-MD induce stabilization of functional Hsp90-MD residues from the catalytic loop (391-SEDLPLNLSREMLQQ-405) (Figure 4C). This cochaperone-mediated stabilization enforces the closed state of the catalytic loop and rigidifies the closed Hsp90 dimer consistent with the functional role of p23 in partially inhibiting the ATPase cycle. The two major distribution peaks of the distance fluctuation stability profile for FKBP51 are aligned with cochaperone-Hsp90A interfaces, where the first cluster corresponds to D72, N74, E75 interacting with Hsp90-MD client binding sites, and the second cluster is formed by conserved residues Y409, M412, and F413 interacting with the Hsp90-CTD helices (residues 658-696) (Figure 4E). In the Hsp90-GR-P23 complex, binding-induced stabilization in the client is particularly strong in the GR pre-helix 1 region (residues 523-531) interacting inside the Hsp90 tunnel, and GR helix 1 ( residues 532–539) stacking against Hsp90-CTD amphipathic helical hairpin (Figure 4F).

To quantify the extent of protection of the Hsp90 residues, we also evaluated the ensemble-based relative solvent accessibilities (RSA) of the protein residues^106^ and examined the scatter plots between root mean square fluctuations (RMSF) and RSA values in the Hsp90-FKBP51-P23 complex (Supporting Information, Figure S1) and in the Hsp90-GR-P23 complex (Supporting Information, Figure S2). The distributions revealed an appreciable degree of correlation between these parameters ( Pearson correlation coefficient R ∼0.55-0.6), highlighting the greater rigidity of the Hsp90 dimer and particularly cochaperone-bound Hsp90-B protomer in the Hsp90-FKBP51-P23 complex ( Supporting Information, Figures S1B). A moderately greater mobility of the Hsp90 residues was seen in the Hsp90-GR-P23 complex, but the level of correlation between the RMSF and RSA values remained the same (Supporting Information, Figures S2). In general, in both complexes Hsp90 residues with significant solvent accessibility (RSA ∼60-70%) could often experience rather moderate fluctuations in the closed dimer conformations, indicating that cochaperone and client binding can curtail the Hsp90 flexibility, which is particularly evident in the Hsp90-FKBP51-P23 complex.

We also analyzed the maps of dynamic cross-residue correlations in the Hsp90 complexes that showed how cochaperones and GR client can affect the long-range couplings in the Hsp90 dimer (Figure 5A-C) . In the Hsp90-P23 complex, we observed the positive correlations of the NTD and CTD regions within the individual protomers and between the M-domains of the two protomers (Figure 5A). At the same time, the cross-correlation maps pointed to the anti-correlated fluctuations between the NTD of one protomer A and the CTD of the protomer B. This is consistent with the experimental data^13^ and our earlier studies of the yeast Hsp82 chaperone^74–77^ showing that anti-correlated motions between the NTDs and CTDs in the Hsp90 dimer are the intrinsic dynamic features of the Hsp90 chaperone and are central to allosteric conformational changes during the chaperone cycle. In the Hsp90-FKBP51-P23 complex, cochaperone binding promoted anti-correlated fluctuations of the NTD and CTD within Hsp90-A protomer, while partial immobilization of the Hp90B motions produced strong positive couplings within the protomer B as well as positive correlations between Hsp90A-CTD and Hsp90-B (Figure 5B). As a result, P23 and FKBP51 binding can curtail coordinated anticorrelated fluctuations of the NTDs and CTDs, thus restraining allosteric motions and arresting the closed Hsp90 dimer conformation. In the Hsp90-GR-P23 maturation complex, moderate but more synchronous anti-correlated couplings between NTD of one protomer and CTD of the other protomer were seen (Figure 5C). These patterns of dynamic changes can be seen in the structural mapping of the Hsp90 mobility along the essential low frequency modes (Figure 5D-F). While Hsp90-P23 and Hsp90-GR-P23 complexes feature coordinated global movements of the NTD and CTD regions (Figure 5D,F), FKBP51 cochaperone together with P23 can partially immobilize Hsp90 motions along slow modes (Figure 5B).

**Figure 5.**
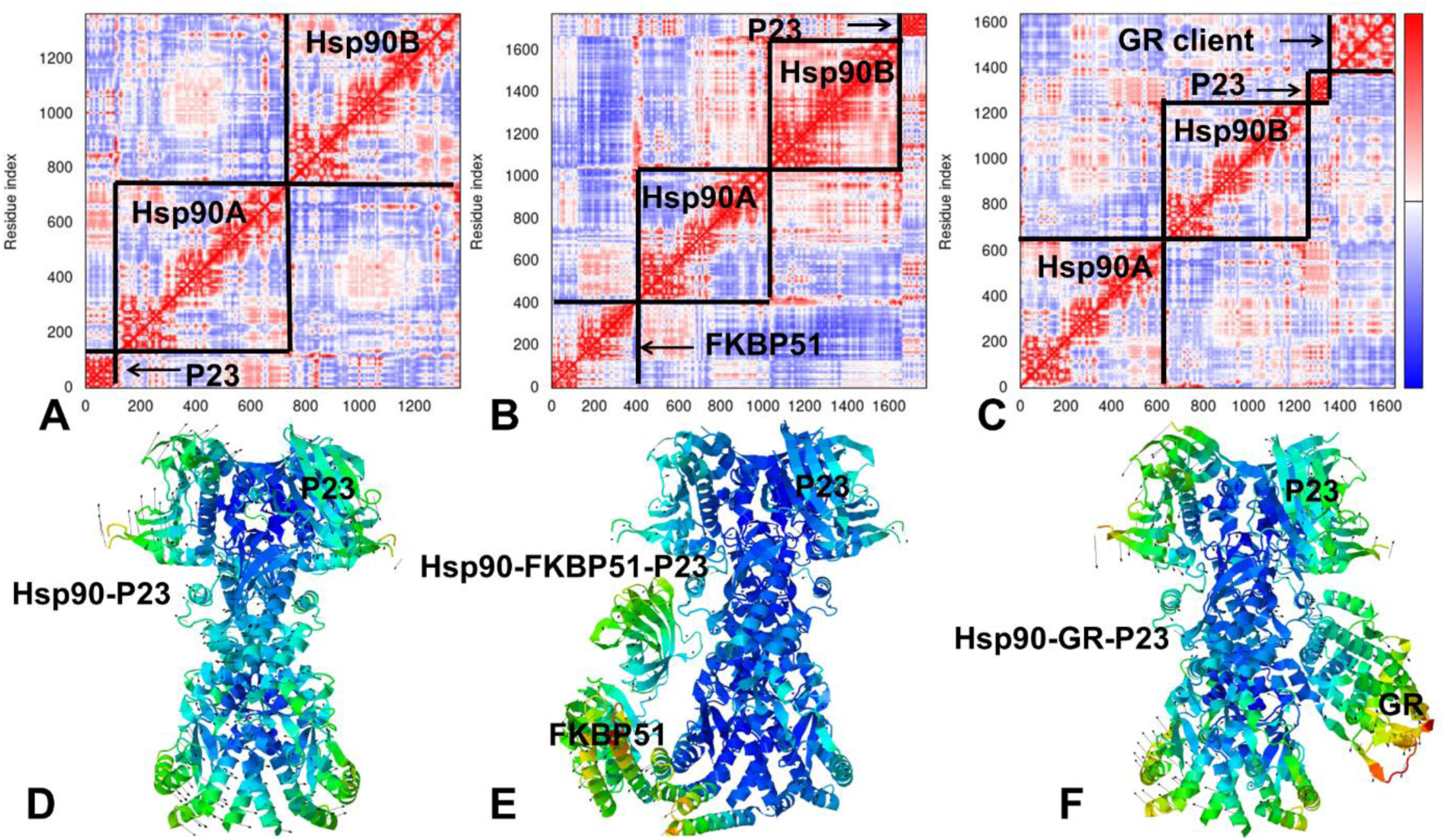
Functional dynamics of the Hsp90 complexes. The covariance maps of dynamic cross-correlations between pairs of residues in the Hsp90-P23 complex (A), Hsp90-FKBP51-P23 complex (B) and Hsp90-GR-P23 complex (C). Cross-correlations of residue-based fluctuations vary between +1 (correlated motion; fluctuation vectors in the same direction, colored in dark red) and -1 (anti-correlated motions; fluctuation vectors in the same direction, colored in dark blue). The values > 0.5 are colored in dark red and the lower bound in the color bar indicates the value of the most anti-correlated pairs. The boxes highlight cross-correlations for the Hsp90 protomers, FKBP51, P23 and GR proteins. Structural maps of correlated motions in the low frequency modes for Hsp90-P23 complex (D), Hsp90-FKBP51-P23 complex (E) and Hsp90-GR-P23 complex (F). The structures are in ribbons with the rigidity-to-flexibility scale colored from blue to red.

GR client protein is also involved in coordinated movements along with Hsp90-NTD and CTD regions. This dynamically controlled allosteric couplings may be necessary for a “client sliding” mechanism proposed in the original structural studies^40,41^ by Agard’s group in which Hsp90 can adjust the client binding site to provide protected refolding of clients before they exit the Hsp90 internal tunnel to become directly stabilized by co-chaperones. The structural map for the Hsp90-FKBP51-P23 complex highlighted how FKBP51 is dynamically anchored to the Hsp90-CTD dimerization helices, while allowing for docking motions of the FK1 domain on the Hsp90-MD client site (Figure 5B). This mode of collective motions may enable FKBP51 to dynamically engage on the Hsp90-MD client site while maintaining the anchoring hinge at the primary recognition site owing to strong FKBP51-TPR interactions with the Hsp90-CTD regulatory helices (Figure 5B). The observed dynamic patterns of local and global motions are consistent with and illustrate the mechanism in which the TPR domain helix functions as a critical specificity element of Hsp90 recognition and at the same time affords the PPIase-active FK1 domain sufficient mobility to properly adjust its position in the Hsp90 client binding site for adaptive client refolding and maturation stages of the cycle.^37^

### Energetic Cartography of the Hsp90 Interactions: Mutational Scanning and Sensitivity Analysis Reveal Structural Stability Centers and Binding Interface Hotspots

To examine the protein stability and binding determinants of the Hsp90 interactions with cochaperones and client protein, we conducted a systematic mutational scanning of protein residues in the Hsp90-FKBP51-P23 (Figure 6) and Hsp90-GR-P23 complexes (Figure 7). Unlike traditional structure-based alanine scanning, we employed the equilibrium ensembles generated by simulations to perform a comprehensive mutational sensitivity analysis of protein residues using statistical averages of the free energy changes. In this analysis, the Hsp90 intermolecular interface residues with FBP51, P23 and GR proteins are defined by the 5 Å contact distance threshold. The interfacial residues for the interacting protein are then systematically mutated to produce mutational sensitivity heatmaps (Figures 6,7). In addition, we separately evaluated contributions of the Hsp90 residues to protein stability of the Hsp90 dimer in the complexes by mutating all chaperone residues and computing the effect of mutations on the inter-protomer interactions and stability of the Hsp90 dimer. Using this mutational cartography approach, we first examined the biding free energy changes in the Hsp90-FKBP51-P23 complex (Figure 6). Consistent with the dynamic signatures of this complex revealing a broadly distributed binding-induced immobilization of the Hsp90 motions, the mutational sensitivity heatmaps of the Hsp90 reflected the extensive network of chaperone interactions featuring a number of intermolecular binding hotpots (Figure 6 A,B). The common binding hotspots in both protomers corresponded to the Hsp90-CTD residues 655-698 involved in the interactions with FKBP51 (Figure 6A,B). In this context, it is notable that coordinated opening and closing of the Hsp90-CTDs and long-range communication between the Hsp90 domains is regulated by these CTD helices.^13^ Moreover, these Hsp90-CTD regions were experimentally confirmed as energetic hotspots of the C-terminal dimerization and Hsp90 stability as mutations in the CTD helices αH17 and αH18 disrupt the dimerization interface and impair chaperone activity.^136^ These findings also supported the experimental hypothesis that FKBP51 primary recognition is enabled by these Hsp90 CTD helices αH17 and αH18 of each protomer.^37^ Indeed, we observed that a range of Hsp90-CTD residues, most notably including the Hsp90-CTD αH18 helix (residues 680-697) can act as mutation-sensitive binding hotspots in both protomers as residue substitutions of R690, M691, L694, and G695 sites induced consistent destabilization changes (Figure 6A,B).

**Figure 6.**
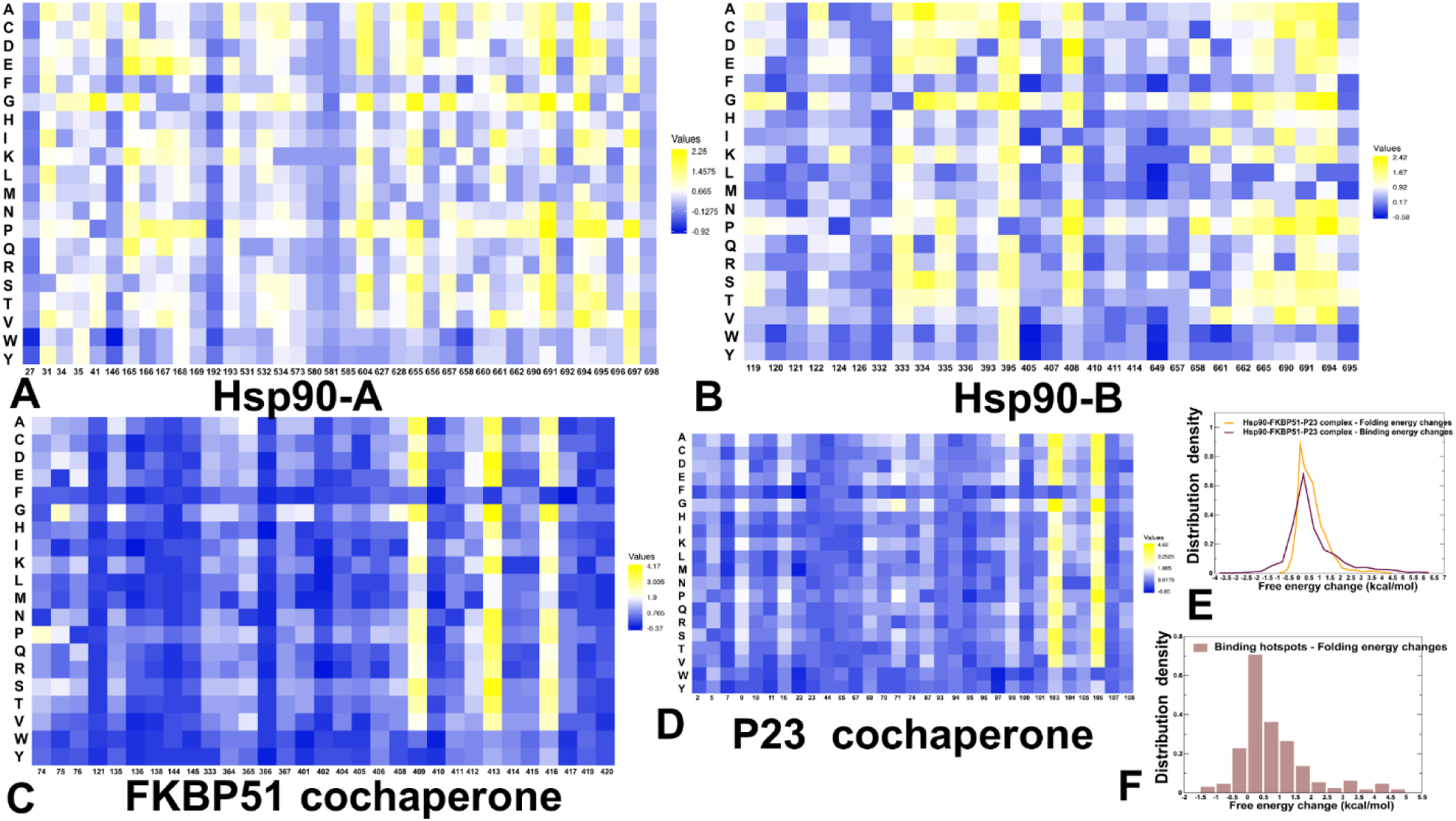
Ensemble-based dynamic mutational profiling of the intermolecular interfaces in the Hsp90-FKBP51-P23 complex. The mutational scanning heatmaps are shown for the Hsp90-A protomer (A), Hsp90-B protomer (B), FKBP51 cochaperone (C), and P23 cochaperone (D). The heatmaps show the computed binding free energy changes for 20 single mutations of the interfacial positions. The cells in the heatmap are colored using a 3-colored scale – from blue to white and to yellow, with yellow indicating the largest unfavorable effect on binding. The standard errors of the mean for binding free energy changes were based on MD trajectories and selected samples from 5 trajectories ( a total of 1,000 samples) are within ∼ 0.11-0.18 kcal/mol using averages over different trajectories. The distribution of the folding free energy changes in the Hsp90 dimer based on mutational scanning of all Hsp90 residues (in orange lines) and biding energy changes based on mutational scanning of all intermolecular residues ( in maroon lines) (E). The distribution of the folding free energy changes in the Hsp90 dimer for the Hsp90 residues involved in the intermolecular binding interfaces (F).

**Figure 7.**
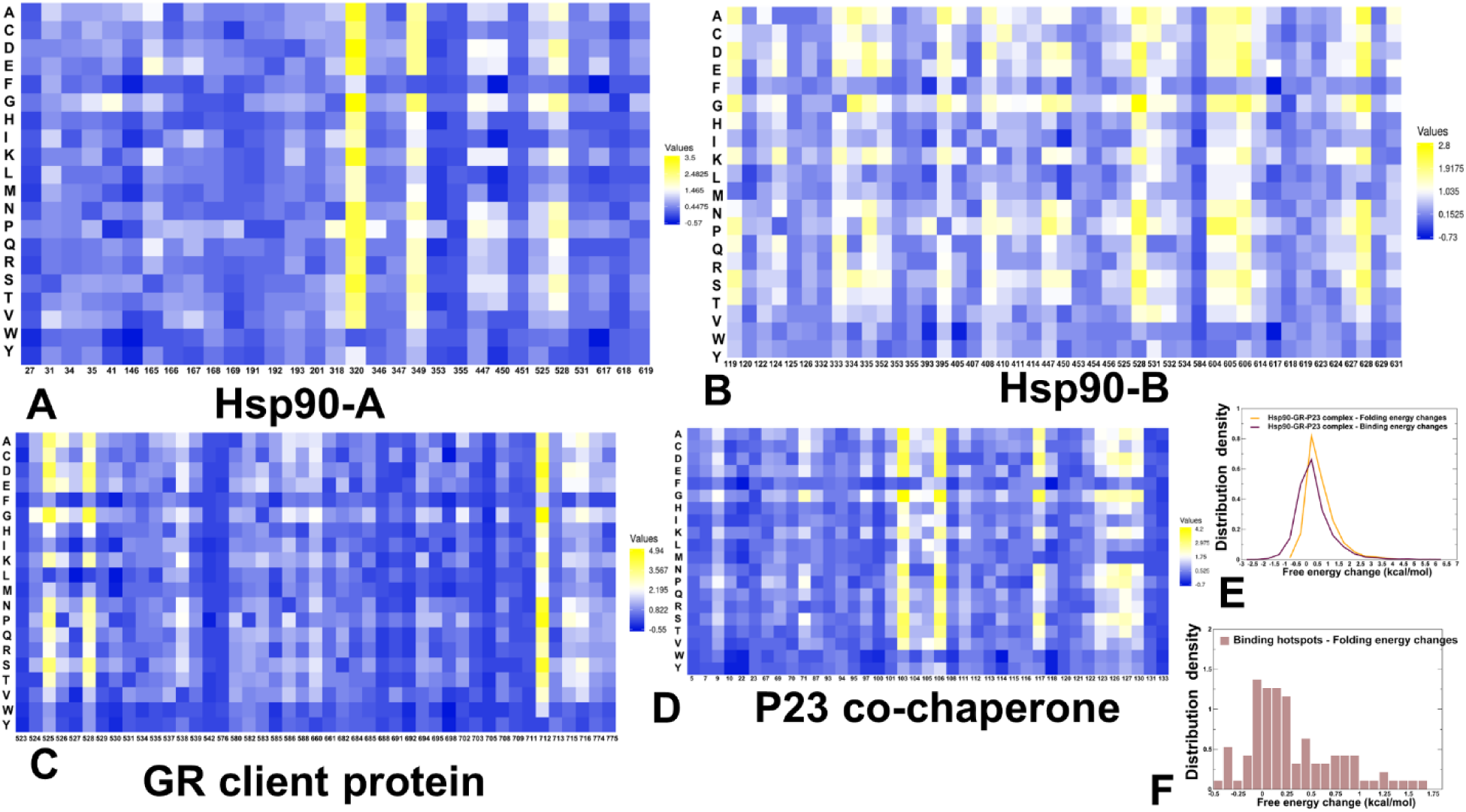
Ensemble-based dynamic mutational profiling of the intermolecular interfaces in the Hsp90-GR-P23 complex. The mutational scanning heatmaps are shown for the Hsp90-A protomer (A), Hsp90-B protomer (B), GR client protein (C), and P23 cochaperone (D). The heatmaps show the computed binding free energy changes for 20 single mutations of the interfacial positions. The cells in the heatmap are colored using a 3-colored scale – from blue to white and to yellow, with yellow indicating the largest unfavorable effect on binding. The standard errors of the mean for binding free energy changes were based on selected samples from 5 separate trajectories ( a total of 1,000 samples) are within ∼ 0.08-0.17 kcal/mol using averages over different trajectories. The distribution of the folding free energy changes in the Hsp90 dimer based on mutational scanning of all Hsp90 residues (in orange lines) and biding energy changes based on mutational scanning of all intermolecular residues ( in maroon lines) (E). The distribution of the folding free energy changes in the Hsp90 dimer for the Hsp90 residues involved in the intermolecular binding interfaces (F).

In reciprocal fashion, the FKBP51 hydrophobic residues I408, Y409, M412, F413, and F416 in the C-terminal extension helix H7e that interact with the Hp90-CTD helices αH17 and αH18 corresponded to the exclusive binding hotspots of the FKBP51 cochaperone (Figure 6C). Consistent with the observed mutational sensitivity of these residues to binding with Hsp90, residues Y409, M412, and F413, are strictly conserved among the immunophilin class of TPR cochaperones.^37^ In agreement with the experiments, ^37^ our results confirmed that these interactions are the primary determinants of conserved Hsp90-specific recognition mechanism for FKBP51 cochaperones. The significance of these findings stems from the fact that FKBP51 binding is driven by targeting the regulatory Hsp90-CTD helices that are also essential for dimerization and responsible for modulation of conformational changes in the Hsp90 chaperone. We argue that as a result, FKBP51 can exploit the localized primary binding hotspots on the Hsp90-CTD switch regions to integrate this cochaperone into allosteric Hsp90 machinery and influence long-range communications required for cochaperone-dependent clients.

The results also pointed to a minor role of the second FKBP51 interacting cluster ( D72, N74 and E75 from the FK1 active site (Figure 2) as these residues appeared to be fairly tolerant to mutational changes and allow for weak dynamic interactions with the Hp90-MD client site (Figure 6C). These findings are also consistent with the collective dynamics of the Hsp90-FKBP51-P23 complex (Figure 5B,E) showing the dynamic nature of the interactions in this binding cluster. We also found that the conserved hydrophobic residues F103 and W106 of p23 are pronounced binding hotspots of the Hsp90-P23 interactions in the complex (Figure 6D). These P23 residues interact with hydrophobic Hsp90-MD sites (L335, L394, P395, I408, and V411). Our results confirmed the mutational sensitivity and importance of this binding interface as a primary recognition site for P23 integration into the Hsp90 machinery. Importantly, the P23 and FKBP51 cochaperones are anchored to the Hsp90 dimer using key regulatory regions across all domains : P23 stabilizes the catalytic loop (391-SEDLPLNLSREMLQQ-405) and the NTD-MD interface; FKBP51 constraints collective motions of the Hsp90-CTD switch helices and modulates dynamics of the Hsp90-MD client site. Hence, cochaperones may exploit several spatially separated hotspots to efficiently control allosteric communications and influence conformational changes in the Hsp90 dimer during the ATPase cycle.

To compare the mutation-induced changes in the protein stability and binding interactions of the Hsp90 dimer, we also performed a systematic mutational scanning of all Hsp90 residues and evaluated the resulting FoldX free energies of the Hsp90 dimer (Figure 6E). The density distributions of mutation-induced folding free energy changes and binding energy changes showed a generally similar profile (Figure 6E). In general, mutations tend to have only marginally stronger effects on the free energy of folding than binding in the Hsp90-FKBP51-P23 complex. Consistent with the deep mutational scanning experiments that mapped the energetic landscapes for several protein binding domains^137^, our results showed that mutations that affect folding free energies are also more numerous and more widely distributed throughout the Hsp90 domains, with many positions sensitive to substitutions (Figure 6E,F). Notably, the distribution of folding free energy changes is entirely determined by the destabilization stability changes (ΔΔG > 0.0) but the average Foldx free energy values peaked near ΔΔG ∼ 0.5 kcal/mol suggesting only minor destabilization effect (Figure 6E). In some contrast, the binding free energy changes have a noticeable density for the negative ΔΔG values corresponding to the increased binding, but the distribution peak ( ΔΔG ∼ 0.2-0.3 kcal/mol) and the bulk of the density showed that mutations generally have a destabilization effect on the intermolecular binding (Figure 6E). Mutations that disrupt binding may often show pleiotropic cooperation by reducing both binding and folding stability^137^, and the distribution of folding energy changes for the binding hotpot sites showed a strong peak for marginally destabilizing ΔΔG ∼ 0.2-0.3 kcal/mol and smaller peaks for appreciably larger ΔΔG ∼ 1.0-2.0 kcal/mol(Figure 6F). This is consistent with the notion that residues involved in the substrate binding or protein binding interfaces are not necessarily optimal for protein stability. Interestingly, these results may be general for a range of protein interaction domains as similar trends were observed in the recent deep mutational scanning analysis of protein binding domains.^137^

In the Hsp90-GR-P23 complex, despite a large binding interface of the Hsp90-A protomer with GR client there are only two pronounced binding hotspots W320 and F349 forming the key intermolecular contacts with the client protein (Figure 7A). Remarkably, W320 also emerged as the strongest binding energy hotspot of the Hsp90-GR interactions in the Hsp90-Hsp70-Hop-GR loading complex.^85^ These findings are consistent with the unique functional role of the Hsp90 W320 (W300 in yeast Hsp82) as a universal molecular switch that is required for Hsp90 regulation during the ATPase cycle^59^ and integration of the Aha1 cochaperone.^61^

The observed high mutational sensitivity of W320 in the complexes with GR client is also consistent with functional experiments showing that modifications in this position can compromise GR binding and activation.^60,138^ In fact, W320 residue was also highlighted as regulatory hotspot in evolutionary analysis of the adaptive potential of the Hsp90-MD showing the important role of this center in stabilization of client-binding interfaces.^138^ Mutational scanning also highlighted the role of F349 residue for binding with the GR client protein (Figure 7A).

The Hsp90 interactions with the unfolded pre-helix 1 region of GR client (residues 523-531) that is inserted inside the Hsp90 tunnel (Figure 3) are dynamic and yielded binding hotspot Y528 in the Hsp90-A (Figure 7A) and hotspot positions Y528 and Q531 in the Hsp90-B protomer (Figure 7B). The Hsp90B-GR interface is formed between a short GR helix 1 (residues 532–539) and the amphipathic helical hairpin on Hsp90-B (residues 607–621) (Figure 3). The observed binding hotpots W606 and M628 in the Hsp90-B (Figure 7B) correspond to conformational switch regions that mediate allosteric communications between NTD and CTD regions. Hence, the Hsp90 binding hotspots for GR client are aligned with the regulatory motifs that are known to mediate allosteric changes in the chaperone.

Common to all complexes, F103 and W106 corresponded to P23 binding hotspots (Figure 7C), while residues L525 and L528 from the unstructured GR pre-helix 1 (residues 523-531) emerged as mutation-sensitive positions on the GR client protein (Figure 7D). The overall distributions of folding free energy changes and intermolecular binding energy changes (Figure 7E) are similar to the ones obtained for Hsp90-FKBP51-P23 complex (Figure 6E). The distribution of folding free energy changes is dominated by destabilization stability changes (ΔΔG > 0.0) with a similar peak near ΔΔG ∼ 0.5 kcal/mol (Figure 7E). The distribution of folding free energy change for the binding hotspots in the Hsp90-GR-P23 complex is extremely broad, peaking near ΔΔG ∼ 0.0-0.5 kcal/mol (Figure 7F). The tail of larger ΔΔG values is associated with stability changes for partly buried Hsp90B residues Y528 and W606 (Figure 3). However, for solvent-exposed binding hotspots on Hsp90 such as W320, mutations can only marginally change protein stability. A similar trend is also seen in the distribution of Hsp90 stability changes in the Hsp90-FKBP51-P23 complex (Figure 6F).

To summarize, results demonstrated several important trends. First, the binding energy hotspots in the Hsp90 dimer are localized in the key regulatory regions, most notably the Hsp90-CTD helical regions responsible for modulation of chaperone dimerization and allosteric communication. Second, we determined that GR client integration into the Hsp90 chaperone system is critically dependent on the interactions with the universal regulatory switch center W320 which also modulates a specific Hp90 conformation needed for client recognition. The central finding of the energetic mutational scanning is that the client- and cochaperone-specific binding interfaces with the Hsp90 induce formation of the binding hotspot centers on Hsp90 that are also associated with important regulatory functions and modulate allosteric communication in the Hsp90 dimer. Hence, by targeting these allosteric regulatory check points, the interacting client proteins can elicit specific conformational changes and induce changes in the global interaction network required to accelerate or slow down progression of the ATPase cycle.

### Perturbation Response Scanning Unveils Allosteric Effectors of Long-Range Communication in the Hsp90 Complexes

Molecular simulations and mutational scanning of binding interactions in the Hsp90 complexes provided a detailed characterization of the binding interfaces and key drivers of binding affinity. Here, we complemented these results with the computational analysis of allosteric interactions in the Hsp90 multiprotein regulatory complexes using perturbation-based profiling approaches that quantify allosteric propensities of protein residues and identify functional allosteric hotspots of residue interaction networks. Here, we adopted the perturbation-response scanning (PRS) approach^139–141^ as implemented by Bahar and colleagues^142,143^ to identify the allosteric effector sites defined as residues where the perturbations cause the largest long-range effect on the dynamics of other residues. In this implementation of PRS, a perturbation force is applied to one residue at a time, and the response of the protein system is measured according to Hooke’s law as a displacement vector **ΔR**(*i*) = **H**^−**1**^**F**(i) that is then translated into *N*×*N* PRS matrix. In this matrix, the *ij*th element evaluates the sensitivity of mode *i* to perturbation at position *j* . By using this approach, we compared the PRS effector profiles for the Hsp90 protein residues that measure the ability of a given residue to influence dynamic changes in other residues in the Hsp90-P23 complex (Figure 8A,D), Hsp90-FKBP51-P23 complex (Figure 8B,E), and Hsp90-GR-P23 assembly (Figure 8C,F). In addition, we also computed the PRS sensor profiles to characterize the propensities of residues to serve as receivers of perturbations in other sites and determine the hotspot positions for main sensors of allosteric perturbations. We test a hypothesis that the regulatory conformational switches in the Hsp90 should exhibit strong allosteric propensities and correspond to the PRS distribution peaks defined as potential effector or sensor hotspots of long-range interactions in the Hsp90 complexes (Figure 8).

**Figure 8.**
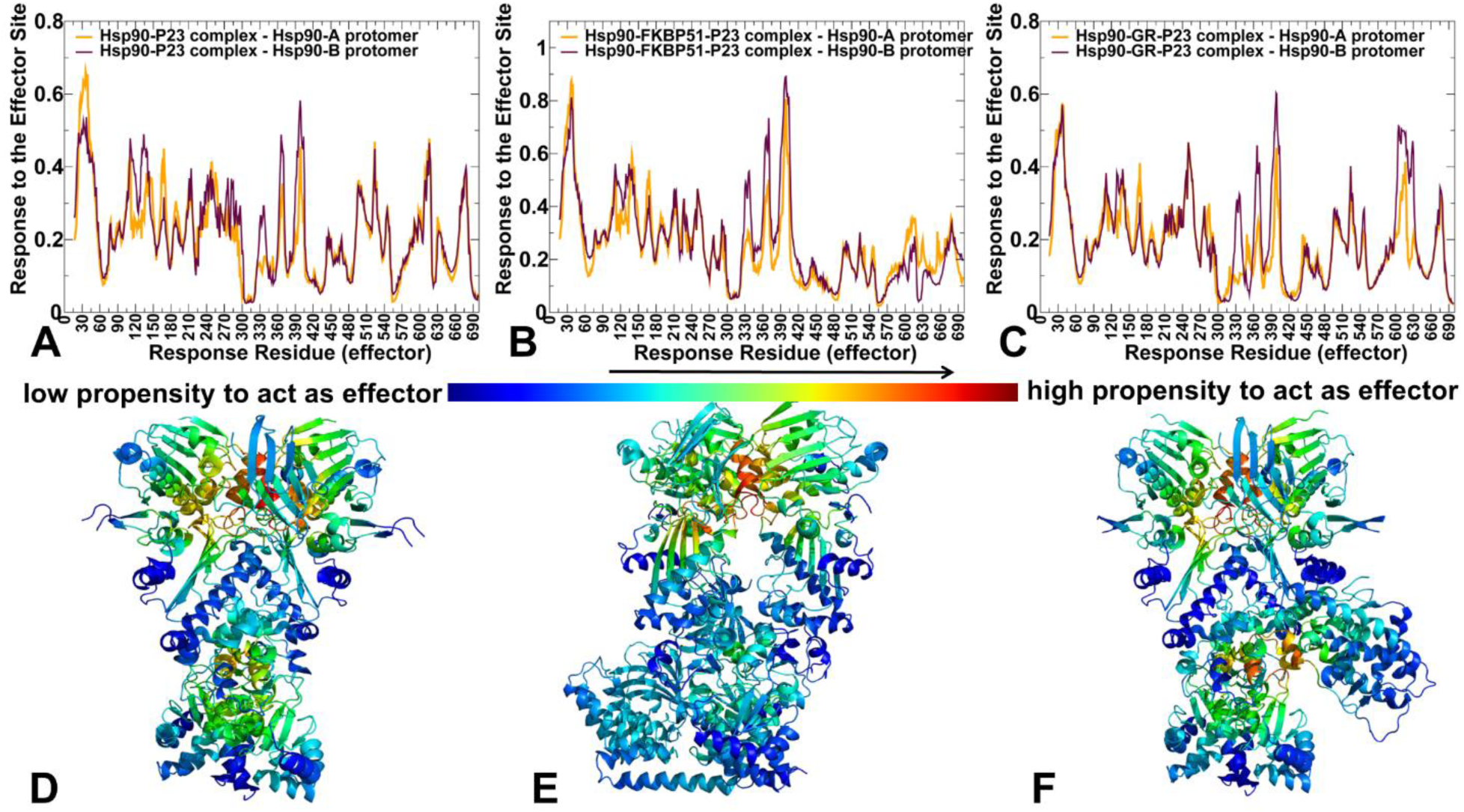
PRS analysis and the effector residue profiles of the Hsp90 regulatory complexes. The residue-based effector profiles of the Hsp90 protomers in the Hsp90-P23 complex (A), Hsp90-FKBP51-P23 complex (B), and Hsp90-GR-P23 complex (C). Hsp90-A protomer is in orange lines, Hsp90-B protomer is in maroon-colored lines. Structural maps of the PRS effector profiles for the Hsp90-P23 complex (D), Hsp90-FKBP51-P23 complex (E), and Hsp90-GR-P23 complex (F). The structure is colored according to the effector potential with blue-to-red color spectrum reflecting the increase of the effector capacity.

The PRS effector analysis provided a quantitative assessment of mediating propensities of functional regions in the Hsp90 complexes (Figure 8). In the Hsp90-P23 complex, the peaks of the effector profile corresponded to the ATP-binding pocket NTD residues (T31,Y33) near the catalytic loop of the Hsp90-MD (residues 391-410) and NTD-MD interfaces (Figure 8A). Functionally important residues near the NTD-MD interface region (V368, F3690, R400, E401, and Q404) include catalytic residues and hydrophobic groups that participate in allosteric communication and stabilize the inter-domain networks required for the formation of active dimer.^74–77^ Although the strongest effector peaks are consolidated in the Hsp90-MD regions, there are two sharp peaks associated with the regulatory Hsp90-CTD helices (residues 658-696) (Figure 8A). Based on this analysis and our energetic analysis, mutations in the allosteric effector regions could have a deleterious effect on the protein stability and allostery, thus severely compromising chaperone activity. In this context, it noteworthy that evolutionary and functional analysis of the Hsp90 showed that beneficial mutations are consolidated in the Hsp90-MD positions 376–386, while deleterious for activity mutations preferentially target client-binding loop (positions 347–361) and residues 396-405 in the Hsp90-MD region.^138^ The PRS analysis identified this Hsp90-MD segment 396-405 as the most prominent effector center (Figure 8A), pointing to a considerable mutational sensitivity of key allosteric hotspots in the otherwise highly adaptive chaperone system. Structural mapping of the PRS effector profile for the Hp90-P23 complex illustrated the distribution of major clusters in the NTD-MD and CTD regions (Figure 8D), showing that the inter-domain regions form allosteric effector centers that can mediate allosteric coupling and communication in the Hsp90 dimer.

In the Hsp90-FKBP51-P23 complex, FKBP51 binding to the Hsp90-MD/CTD regions inflicts a significant redistribution of the effector centers (Figure 8B,E). We observed narrowing of the Hsp90-NTD effector density while the increased peak in the Hsp90-MD region 396-405 that is the conserved effector center in all three complexes. The FKBP51 interactions with the CTD regulatory helices imposed significant constraints on the CTD movements and, as a result, the effector density in these regions is significantly reduced (Figure 8B). Structural map of the effector distribution highlighted these changes, pointing out to effector centers localized in the NTD-MD region (Figure 8E). Hence, FKBP51 binding may partly impair the inter-domain dynamic allostery in the Hsp90 and arrest the closed Hsp90 conformation, allowing for local modulation of flexibility at the FK1 PPIase site interacting with the Hsp90-MD client regions. We argue that such modulation of dynamic allostery may be required to curtail large, coordinated movements in the Hsp90 dimer and ensure only localized dynamic modulation of the Hsp90-MD client site interactions for optimal loading and processing of FKBP51-specific clients.^37^

The PRS effector distribution in the Hsp90-GR-P23 complex (Figure 8C,F) is highly similar to the Hsp90-P23 profile, revealing three distinct clusters of the effector centers in the NTD, MD, and CTD domains. While the Hsp90-NTD and NTD-MD effector clusters are mostly preserved, the effector peaks are particularly prominent near Hsp90-MD amphipathic loop (residues 349–359) and Hsp90-CTD amphipathic helical hairpin (residues 607–621 (Figure 8C). These findings are consistent with the known allosteric role of these regions in mediating dimerization and allosteric communications between NTD and CTD regions.^13,14^ Structural projection of the effector profile highlighted the localization of the effector hotspots near NTD-MD and MD-CTD interdomain interfaces (Figure 8F). Moreover, we noticed that the GR client interface is also enriched by effector sites and involved in dynamic couplings with Hsp90-MD/CTD regions. It is possible that the observed “restoration” of the inter-domain allostery in the Hsp90-GR-P23 complex may be relevant for a “client sliding” mechanism^40,41^ in which Hsp90 fluctuates to facilitate client refolding and sliding to a ligand-bound conformation that is stabilized by both Hsp90 and the p23 tail.

We analyzed the relationship between PRS effector propensities and the relative solvent accessibility (RSA) of the Hsp90 residues (Supporting Information, Figure S3). The correlation scatter plots for the Hsp90 protomers in the Hsp90-FKBP51-P23 complex (Supporting Information, Figure S3A,B) and Hsp90-GR-P23 complex (Supporting Information, Figure S3C,D) are quite similar. We observed only a moderate relationship between these parameters, showing that the Hsp90 residues with the high effector potential in the MD regions are often buried and feature RSA < 20%. This is the case for the Hsp90-MD segment 396-405 acting as the most prominent effector center (Figure 8A) with mutational sensitivity of key allosteric hotspots. At the same time, many allosterically important sites with high PRS effector values are not limited to rigid Hsp90 positions insulated from solvent and may be flexible with the RSAs covering a broad spectrum of solvent accessibilities (Supporting Information, Figure S3). Hence, allosteric effector hotspots in the Hsp90 could often retain a certain degree of mobility, indicative of long-range dynamic correlations in the Hsp90 complexes, where local perturbations in the effector switch centers can induce a cascade of long-range dynamic changes in the system.

The PRS sensor distributions in all three studied complexes were also examined in which profile peaks point to Hsp90 positions extremely sensitive to the perturbations in other sites and typically associated with the moving regions during allosteric changes (Supporting Information, Figure S4). Here, we also observed a similar shape of the PRS profiles for the Hsp90-P23 complex (Supporting Information, Figure S4A,D) and Hsp90-GR-P23 (Supporting Information, Figure S4C,F). As expected, the sensor peak clusters in these complexes are localized in the peripheral areas of the NTD and CTD domains, indicating that these regions would experience coordinated movements which is a signature of the chaperone allostery. Noteworthy, structural mapping of PRS sensor profiles for the Hsp90-GR-P23 complex revealed that GR client regions can undergo concerted dynamic changes along with the peripheral Hsp90-CTD regions (Supporting Information, Figure S4F). The P23 tail helix which is anchored to GR client forms a local hinge that facilitates the activation of the GR by stabilizing dynamic GR helix 12 in its agonist-binding conformation.^41^ The observed “sensing” of the GR regions to allosteric signals and coordinated movements of the GR ligand binding regions with Hsp90-CTDs in the maturation Hsp90-GR-P23 complex may facilitate access to the Hsp70 and be relevant in the established mechanism of Hsp70-mediated unfolding/ligand release and Hsp90-mediated refolding/ligand binding that facilitates rapid responses of the GR protein to changing hormone levels.^144,145^ A different distribution of sensor residues was found for the Hsp90-FKBP51-P23 complex (Supporting Information, Figure S4B,E). In this Hsp90 assembly, the movements of the Hsp90-MD, MD-CTD and Hsp90-CTD regions are largely suppressed, pointing to the mobile receiver regions in the Hsp90-NTD and particularly near the secondary binding site of the FKBP51 ( D72, N74 and E75 from the FK1 active site) (Supporting Information, Figure S4E). These findings are also consistent with the observed dynamic nature of the FKBP51 interactions near the Hsp90-MD client binding site which could allow for engagement different client positions on the Hsp90.

There was no correlation between the PRS sensor propensities and the RSA parameters for the Hsp90 residues (Supporting Information, Figure S5). Indeed, the Hsp90 sensing sites involved in allosterically triggered motions can belong to the inter-domain regions where they are partly buried ( RSA < 40-50%) but also correspond to the peripheral solvent-exposed residues on the NTD and CTD (RSA > 50%). In general, for the structurally adaptive Hsp90 homodimer, the distributions of allosteric effector and receiver centers are weakly correlated with the solvent accessibility and are determined by the intrinsic topology of the Hsp90 fold.

### Allosteric Mutational Profiling of the Residue Interaction Networks in the Hsp90 Complexes Identifies Allosteric Regulatory Switches of the Chaperone Function

By leveraging the extensive dynamic and energetic analysis of the intermolecular interactions in the Hsp90 complexes, we performed dynamic network analysis of the conformational ensembles and introduce a perturbation-based network approach for systematic mutational scanning of allosteric residue preferences and mapping allosteric landscapes. The objective of this analysis was to specify allosteric functional roles of the intermolecular interfaces and identify allosteric regulatory switch centers in the chaperone complexes. By analogy with protein-protein interaction networks, allosteric switches are defined as protein residues that participate in significant edgetic perturbations of the residue interaction network, i.e., sites where mutations disrupt the network connectivity and cause a significant impairment of global network communications.

Using the dynamic network model of residue interactions in which network edges between nodes are weighted using both dynamic correlations^121^ and coevolutionary residue couplings^122^ we computed the ensemble-averaged distributions of several residue-based topological network metrics. The short path residue centrality (SPC) is used to analyze modularity and community organization of the dynamic residue interaction networks. The SPC distributions reflect the extent of residue connectivity in the interaction networks and allow for characterization of the mediating clusters and local communities in the Hsp90 complexes. The Z-score betweenness centrality is based on computing the average shortest path length (ASPL). By systematically introducing mutational changes in the protein positions and using the equilibrium ensemble of the native system, we reevaluate dynamic inter-residue couplings and compute mutation-induced changes in the ASPL parameter. These changes are then averaged over all substitutions in a given residue. In this manner, we characterize the average mutational effect of each node on changes in the network modularity and efficiency of allosteric communications. By identifying residues where mutations induce a significant change in the ASPL metric, we locate allosteric regulatory switches that determine the efficiency of signal transmission in the protein.

The SPC distribution for Hsp90-FKBP51-P23 complex revealed densely populated clusters of high centralities in the Hsp90-MD and CTD regions that are often aligned with the experimentally known functional positions (Figure 9A). A good correspondence was found between the intermolecular binding energy hotspots and the peaks of the SPC distributions. A close inspection of the SPC profiles for the Hsp90 protomers also pointed to the peaks in the Hsp90-NTD dimerization interface formed through interactions of the helix 1 (residues 23-36) and the lid motif (residues 105-137) on both protomers. The functionally important for chaperone activity residues S31, L32, T36 on the Hsp90-NTD as well as residues F118 and M130 of the lid motif corresponded to the centrality peaks (Figure 9A). The high centrality residues are broadly consolidated in the Hsp90-MD regions (residues 380-420), particularly showing high density near 396-LNLSREMLQQ-405 loop (Figure 9A). Remarkably, the SPC centrality distribution for the Hsp90 protomers is very similar to the residue distribution enriched with deleterious mutations in the Hsp90 found in evolutionary studies^138^ and showing a broad peak for the same region in the yeast Hsp90-MD. Accordingly, the high centrality Hsp90-MD residues may be sensitive to mutational perturbations suggesting that deleterious effects of mutations are associated with their mediating role in the residue interaction network. The strong SPC peaks were also seen in the Hsp90-CTD helices (residues 658-696) that are key functional regions that mediate dimerization and allosteric communications between NTD and CTD regions.

**Figure 9.**
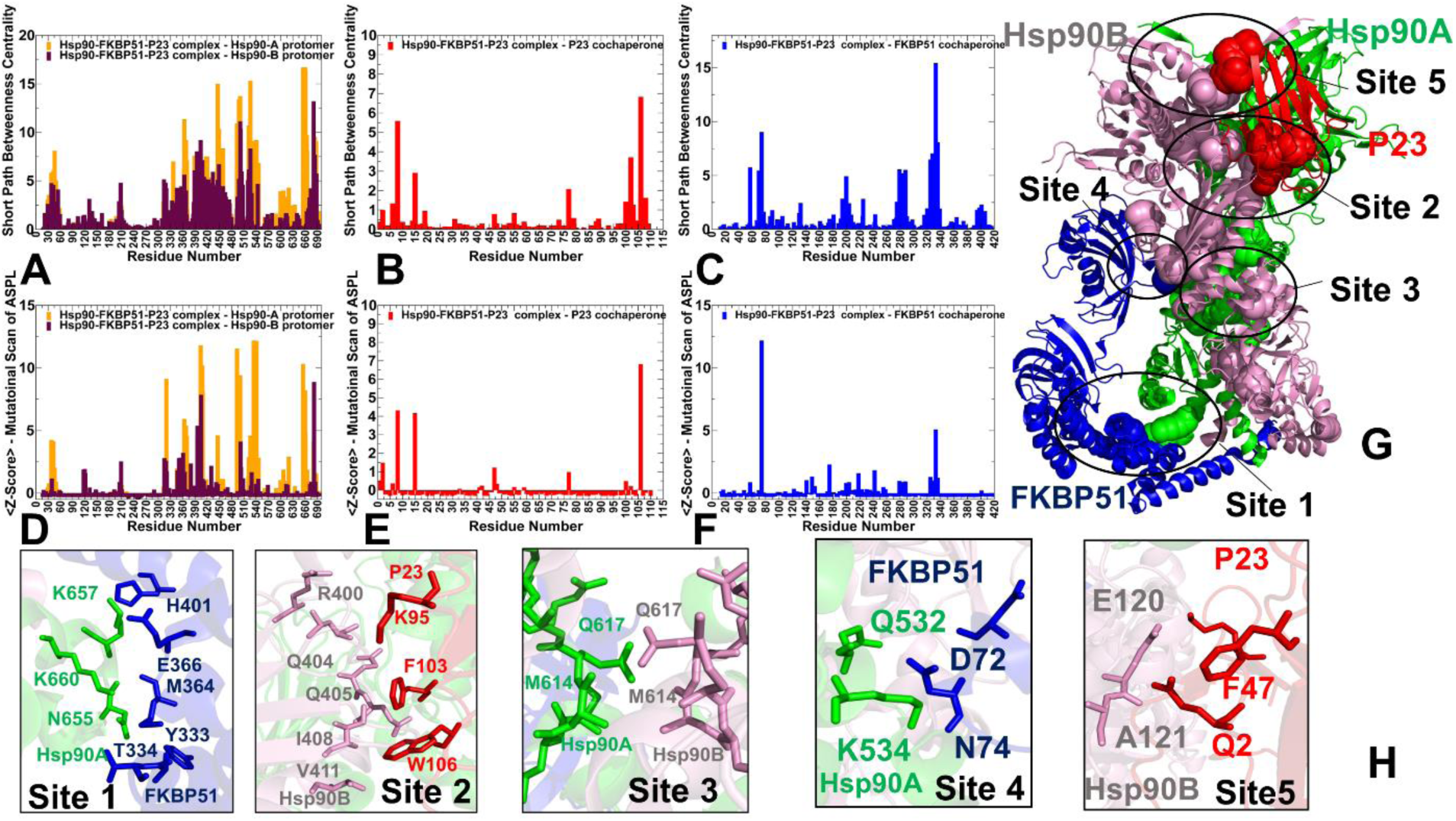
Dynamic network centrality analysis and allosteric mutational profiling of the residue interaction networks in the Hsp90-FKBP51-P23 complex. The residue-based short path betweenness centrality (SPC) profiles for the Hsp90 protomers (A), P23 cochaperone (B) and FKBP51 cochaperone (C). The average mutation-induced changes in the Z-score ASPL parameter based on systematic mutational scanning of allosteric propensities for the Hsp90 protomers (D), P23 cochaperone (E) and FKBP51 cochaperone (F). The SPC and Z-ASPL distributions for the Hsp90-A protomer are in orange lines and Hsp90-B protomer in maroon lines. The profiles for P23 are in red lines and for FKBP51 are in blue lines. Structural mapping of allosteric switch clusters (G). The Hsp90-A in green ribbons, Hsp90-B in pink ribbons, P23 is in red ribbons, and FKBP51 is in blue ribbons. The close-up and atomistic details for major allosteric cluster centers annotated as site 1-site 5 (H). The protein residues involved in these regulatory clusters are shown in sticks and annotated (Hsp90-A in green, Hsp90-B in pink, P23 in red, and FKBP51 in blue).

The SPC distribution for P23 cochaperone highlighted peaks for the conserved F103 and W106 of p23 interacting with Hsp90B-MD (Figure 9B). Interestingly, the SPC profile for the FKBP51 cochaperone displayed a peak corresponding to the helix connecting TPR and FK2 domains ( residues 332-346) and FK1 interfacial residues with Hsp90B-MD (residues 70-80) (Figure 9C). Notably, the primary binding hotspot of FKBP51 with Hsp90 (residues 409-416) featured as only exceedingly small peak, indicating its peripheral role in mediating network connectivity and the long-range communication.

Perturbation-based mutational scanning of allosteric residue propensities in the Hsp90-FKBP1-P23 complex provided information about potential regulatory switches by mapping a space of network-altering allosteric ‘edgetic’ variant sites (Figure 9D-F). The peaks of this profile corresponded to sites where mutational changes lead to dramatic loss in the efficiency of signal transmission as measured by average changes in the ASPL. Importantly, the Z-ASPL distributions displayed a considerable specificity of the peaks, reflecting the unique role of mutation-sensitive network switches (Figure 9D).

Allosteric mutational profiling of the Hsp90-FKBP51-P23 complex revealed five major regulatory clusters that could serve as switches of allosteric communications in the system (Figure 9G, H). Our results showed that the FKP51 helix (residues 350-3640) forms an important switch site at the interface with the Hsp90A-CTD dimerization helix (residues 656-671 (site 1) (Figure 9G,H). This center is coupled with another dynamic switch cluster where FKBP51 residues D72, N74, E75 interact with Hsp90-MD client binding site (site 4) (Figure 9H). We argue that these allosteric hotspots would allow for cochaperone-induced modulation between CTD dimerization helices and Hsp90-MD client sites, thus imposing control on allosteric signaling in the Hsp90 dimer. Another allosteric cluster is formed by the Hsp90-MD residues Q404, Q405, I408 and V411 that interact with the P23 hotspots F103 and W106 (site 2) (Figure 9D,E,H). The corresponding interfacial cluster between P23 and Hsp90B is evolutionary conserved and also corresponded to the binding energy hotspots. According to our results, this regulatory center is also indispensable for allosteric communications in the complex. Indeed, mutations of these residues lead to significant deleterious consequences and compromise Hsp90 regulatory activity and allostery-driven progression of the ATPase cycle. Mutations of corresponding conserved positions in yeast Hsp90 (F121 and W124) tested their effects on the Hsp90 ATPase, showing that F121A/W124A double mutation led to a complete loss of p23’s ability to inhibit Hsp90 and establishing that p23 F121 is essential for the inhibition of the Hsp90 ATPase cycle whereas W124 also contributes to the inhibitory potential.^21^ Finally, allosteric profiling revealed that the intermolecular cluster between Hsp90-CTD amphipathic helical hairpins (residues 607–621) of both protomers (site 3) is a key allosteric hotspot an may coordinate communication with other regulatory clusters (Figure 9G,H). These findings are consistent with the known functional role of these Hsp90-CTD regions in mediating dimerization and allosteric communications between NTD and CTD regions.^13,14^

Allosteric network profiling in the Hsp90-GR-P23 complex (Figure 10) revealed conservation of major network mediating clusters and key allosteric switch centers in the complex. The SPC distribution for the Hsp90 protomers (Figure 10A) and P23 cochaperone (Figure 10B) showed a remarkably similar pattern to the one observed in the Hsp90-FKBP51-P23 complex (Figure 9A,B). The conservation of high centrality clusters in the Hsp90-MD regions (residues 380-420) in both regulatory complexes underscored their evolutionary role as mutations in this MD region are largely deleterious.^138^ These findings further reinforced the notion that the high network centrality of these Hsp90-MD residues makes them sensitive to mutational changes that could lead to drastic alterations in the network connectivity of the Hsp90 chaperone. The SPC distributions for P23 cochaperone in the Hsp90-FKBP51-P23 and Hsp90-GR-P23 complexes are remarkably similar, revealing consistent peaks for the conserved F103 and W106 of p23 interacting with Hsp90B-MD (Figure 10B). The centrality profile of the GR client protein featured 3 broad peak clusters (residues 525-540, 570-590 and 680-700) that are involved in mediating interactions with the Hsp90-MD and CTD regions (Figure 10C).

**Figure 10.**
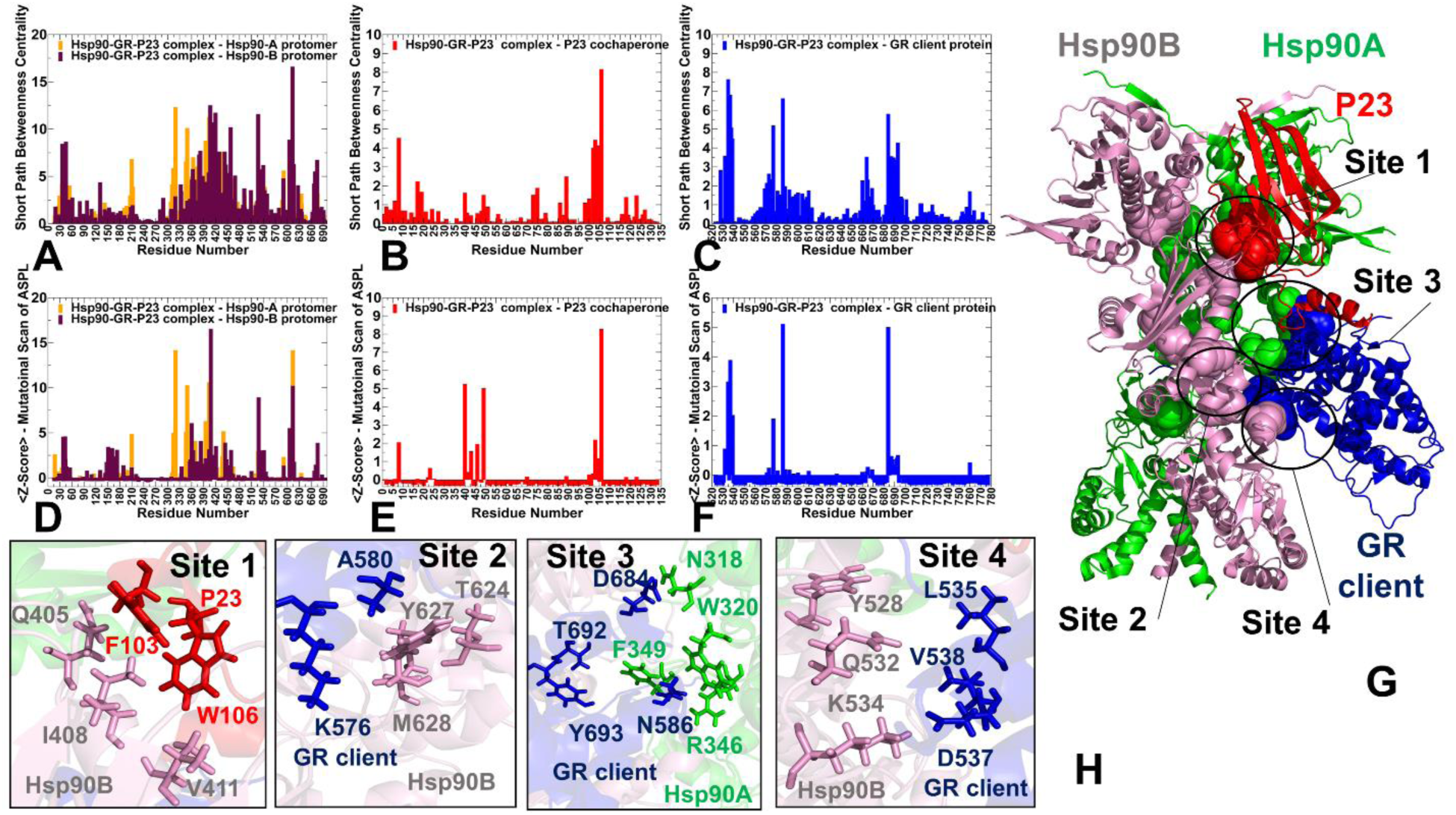
Dynamic network centrality analysis and allosteric mutational profiling of the residue interaction networks in the Hsp90-GR-P23 complex. The residue-based short path betweenness centrality (SPC) profiles for the Hsp90 protomers (A), P23 cochaperone (B) and GR client protein (C). The average mutation-induced changes in the Z-score ASPL parameter based on systematic mutational scanning of allosteric propensities for the Hsp90 protomers (D), P23 cochaperone (E) and GR client protein (F). The SPC and Z-ASPL distributions for the Hsp90-A protomer are in orange lines and Hsp90-B protomer in maroon lines. The profiles for P23 are in red lines and for GR client are in blue lines. Structural mapping of allosteric switch clusters (G). The Hsp90-A in green ribbons, Hsp90-B in pink ribbons, P23 is in red ribbons, and GR protein is in blue ribbons. The close-up and atomistic details for major allosteric cluster centers annotated as site 1-site 5 (H). The protein residues involved in these regulatory clusters are shown in sticks and annotated (Hsp90-A in green, Hsp90-B in pink, P23 in red, and GR client protein in blue).

The Z-ASPL distributions provided a more “granular” characterization of the key allosteric switch centers (Figure 10D-F), displaying well-defined sharp peaks which allowed for a detailed analysis. The pronounced peaks of the Z-ASPL profile are localized in the Hsp90-MD and CTD regions and aligned precisely with the W320 conformational switch, a group of MD residues (L404, I408, L421) in the NTD-MD inter-domain region involved in the P23 interface, as well as CTD helix 607-TANMERIMK-615 (Figure 10D). In particular, W604, A608, M610 and I613 positions in this helix displayed the highest Z-score ASPL values, suggesting that mutations in these CTD sites can lead to dramatic impairment of the allostery in the Hsp90-GR-P23 complex. These findings are an excellent agreement with the experimental data as our analysis singled out the universal regulatory switch Hsp90-MD/W320^59,61^ and identified the Hsp90-CTD helix (residues 607-615) which is experimentally confirmed as a key allosteric switch region that controls Hsp90 conformational changes and can function as a discriminator of selective client binding.^42^ Allosteric profiling of the Hsp90-GR-P23 complex unveiled four clusters that may function as regulatory switch centers. Common to all complexes, the P23 interface between F103/W106 residues and Hsp90B-MD region (residues Q405, I408, V411) forms a conserved allosteric center (site 1) (Figure 10G,H). Interestingly, other three allosteric clusters involve a coordinated group of GR client interaction regions. The key allosteric hotspot is formed by the Hsp90A-MD sites F349/W320 with the GR client (residues N586, H588, K681, S682) (site 3) (Figure 10G,H). The interaction site of the GR pre-helix and helix 1 (residues L535, D537, L538) is coupled with the Y528, Q531 and Q532 positions in the Hsp90-B protomer and the amphipathic helical hairpin region (Y604, G605, W606) (site 4) that form another allosteric switch as evident by sharp Z-ASPL peaks for these positions (Figure 10G,H). Accordingly, the integration of GR client into the Hsp90-P23 system and client-induced modulation of allosteric interaction networks can be orchestrated via these regulatory switches. Our analysis suggested that mutations in these positions could lead to a significant alteration of communication paths in the interaction network. These findings are in good agreement with the experimental data that demonstrated the vital role of yeast Hsp82 CTD resides W585 and M589 ( W604 and M610 in the human Hsp90) for client binding and long-range communication between Hsp90.^42,135^ It is worth stressing that GR client can elicit modulation of the Hsp90 chaperone by targeting the communication switches that are intrinsically present in the apo Hsp90 system.

The mutational scanning maps of the intermolecular interfaces showed that only W320 is a pronounced binding energy hotspot on Hsp90-A protomer, while mutations of other residues with high allosteric potential in the Hsp90-MD and Hsp90-CTD amphipathic helix produced only moderate destabilization changes that are comparable to other interfacial positions. These results suggested that the allosteric centers may not necessarily be the main contributors of protein stability and binding affinity, and their functional role is primarily determined by their topological features and strategic positions at the inter-domain interfaces and client-sensitive switch regions.

## Conclusions

In this study, by combining atomistic simulations, perturbation-based approaches and dynamic network modeling, we carried out a comprehensive profiling of the Hsp90 binding and allosteric interaction networks for the Hsp90-P23, Hsp90-FKBP51-P23 and Hsp90-GR-P23 maturation complexes. The ensemble-based distance fluctuation analysis quantified the effect of cochaperone P23 and GR client on the Hsp90 dynamics, showing that protein binding can curtail coordinated movements of the Hsp90 dimer and slow down the inter-domain allosteric signals, which consistent with the functional role of P23 in arresting the ATPase cycle and stabilizing the closed dimer state. The observed dynamic patterns of global motions in the Hsp90-FKBP51-P23 complex rationalized a mechanism in which the TPR domain helix is a critical specificity element of Hsp90 recognition and allows for dynamic modulation of the FKBP51-Hsp90 interfaces to accommodate cochaperone-specific client binding. Using systematic mutational scanning of protein residues and mutational heatmaps, we characterized the hotspots of protein stability and binding affinity in the Hsp90 complexes, particularly showing that a single W320 switch is the most important binding hotspot in the Hsp90-GR-P23 complex. The central finding of the mutational scanning analysis is that the client- and cochaperone-specific binding interfaces with the Hsp90 involve important regulatory centers that modulate allosteric communication in the Hsp90 dimer. Using perturbation-based network approach for mutational scanning of allosteric residue preferences, this study identified key regulatory centers and characterized allosteric switch clusters that control mechanism of cochaperone-dependent client recognition and remodeling by the Hsp90 chaperone. The results revealed a conserved network of allosteric switches in the Hsp90 complexes, suggesting a mechanism in which P23 and GR proteins are integrated into Hsp90 system by anchoring to the conformational switch points in the Hsp90-MD and Hsp90-CTD regions. In the proposed mechanistic model of the Hsp90 regulation, the general topography of the Hsp90 interaction network and allosteric centers are intrinsically predetermined by the dimer chaperone architecture, whereas activation of specific regulatory switches could be triggered by integration of cochaperones and client proteins near control points of the Hsp90 allostery. The results of this study suggested that the modular organization and allosteric cross-talk between allosteric regulatory clusters in the Hsp90 chaperone can be efficiently exploited by the interacting cochaperones to adapt the Hsp90 allostery and control progression of the ATPase cycle in a client-specific manner.

## SUPPORTING INFORMATION

Figure S1 describes the relationships between the RMSF and RSA values for the Hsp90 protomers, FKBP51 cochaperone, and P23 cochaperone in the Hsp90-FKBP51-P23 complex. Figure S2 illustrates the relationships between the RMSF and RSA values for the Hsp90 protomers, GR client protein, and P23 cochaperone in the Hsp90-GR-P23 complex. Figure S3 describes the correlation analysis between PRS effector residue propensities and the RSA values for the Hsp90 residues in the Hsp90-FKBP51-P23 and Hsp90-GR-P23 complexes. Figure S4 describes the PRS sensor profiles for the Hsp90 protomers in the Hsp90-P23, Hsp90-FKBP51-P23, and Hsp90-GR-P23 regulatory complexes. Figure S5 examines correspondence between the PRS sensor propensities and the RSA parameters for the Hsp90 residues in the in the Hsp90-FKBP51-P23 complex and Hsp90-GR-P23 complexes. This material is available free of charge via the Internet at http://pubs.acs.org.

## AUTHOR INFORMATION

The authors declare no competing financial interest.

## Acknowledgment

The author acknowledges support by the Kay Family Foundation Grant A20-0032.

## For Table of Contents Use Only

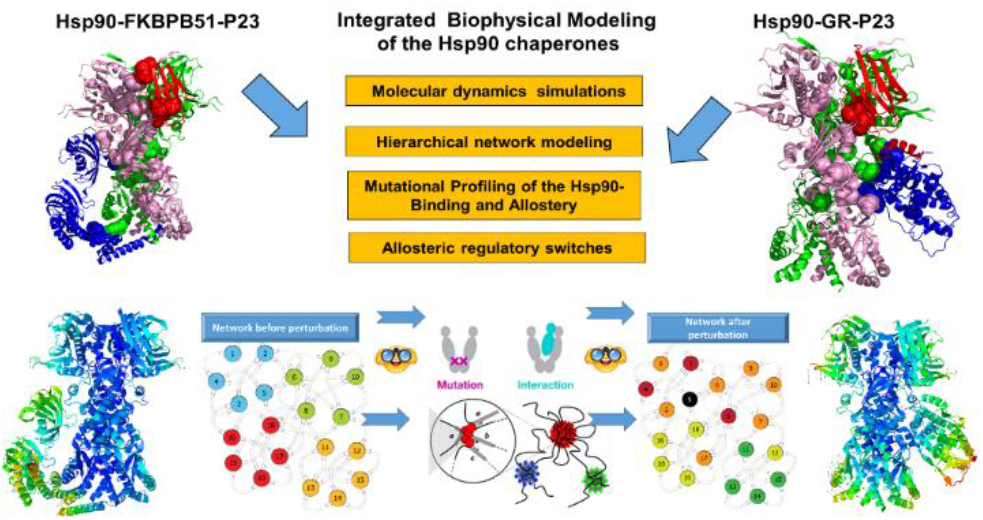

