## Supporting Figures S1-S5 for "Conformational Dynamics and Mechanisms of Client Protein Integration into the Hsp90 Chaperone Controlled by Allosteric Interactions of Regulatory Switches: Perturbation-Based Network Approach for Mutational Profiling of the Hsp90 Binding and Allostery"

Gennady M. Verkhivker[1,2 ‡]

[1]Keck Center for Science and Engineering, Schmid College of Science and Technology, Chapman University, One University Drive, Orange, CA 92866, USA

[2] Depatment of Biomedical and Pharmaceutical Sciences, Chapman University School of Pharmacy, Irvine, CA 92618, USA

‡corresponding author

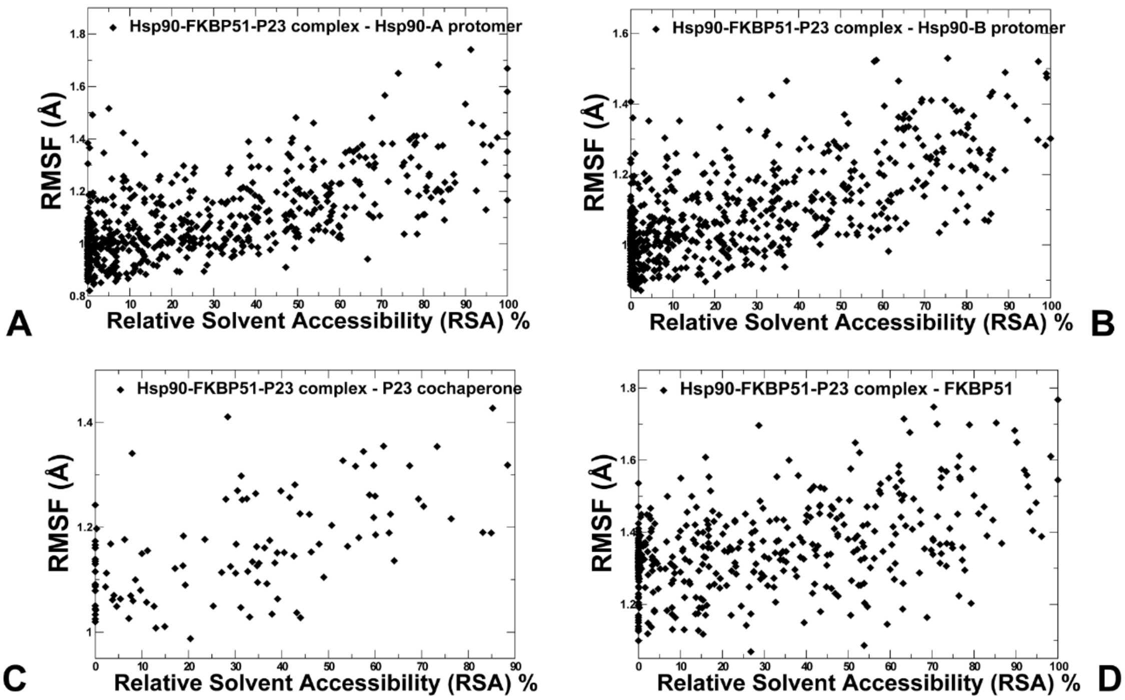


**Figure S1.** The scatter graphs between the RMSF values and the RSA parameters for the protein residues in the Hsp90-FKBP51-P23 complex are shown for Hsp90-A (A), Hsp90-B (B), P23 cochaperone (C) and FKBP51 (D). The residue-base data points are shown in black-colored filled diamonds.

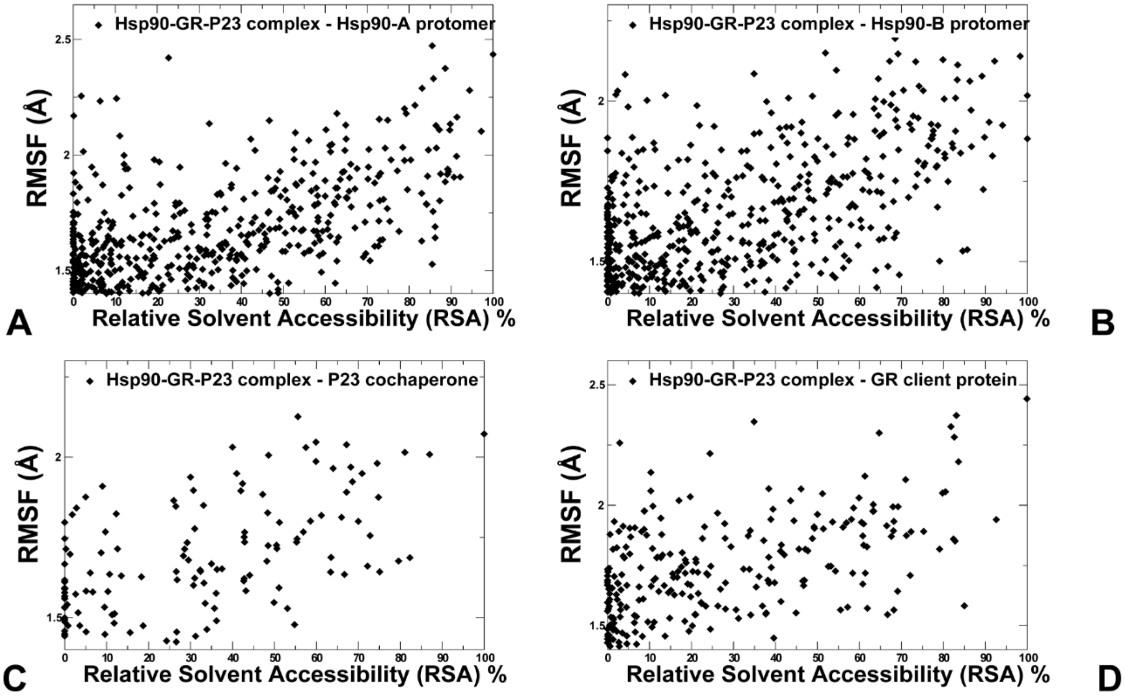


**Figure S2.** The scatter graphs between the RMSF values and the RSA parameters for the protein residues in the Hsp90-GR-P23 complex are shown for Hsp90-A (A), Hsp90-B (B), P23 cochaperone (C) and GR client protein (D). The residue-base data points are shown in black-colored filled diamonds.

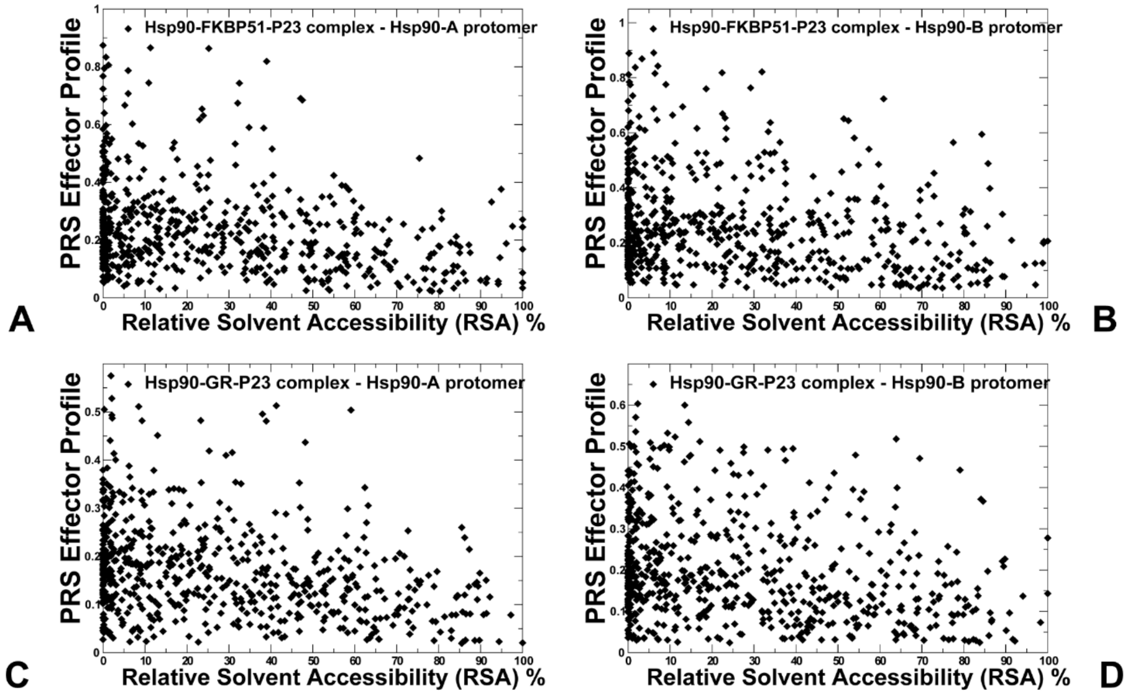


**Figure S3.** The scatter graphs between the PRS effector profile values and the RSA parameters for the Hsp90 protomer residues in the Hsp90-FKBP51-P23 complex (A,B) and Hsp90-GR-P23 complex (C,D). The residue-base data points are shown in black-colored filled diamonds.

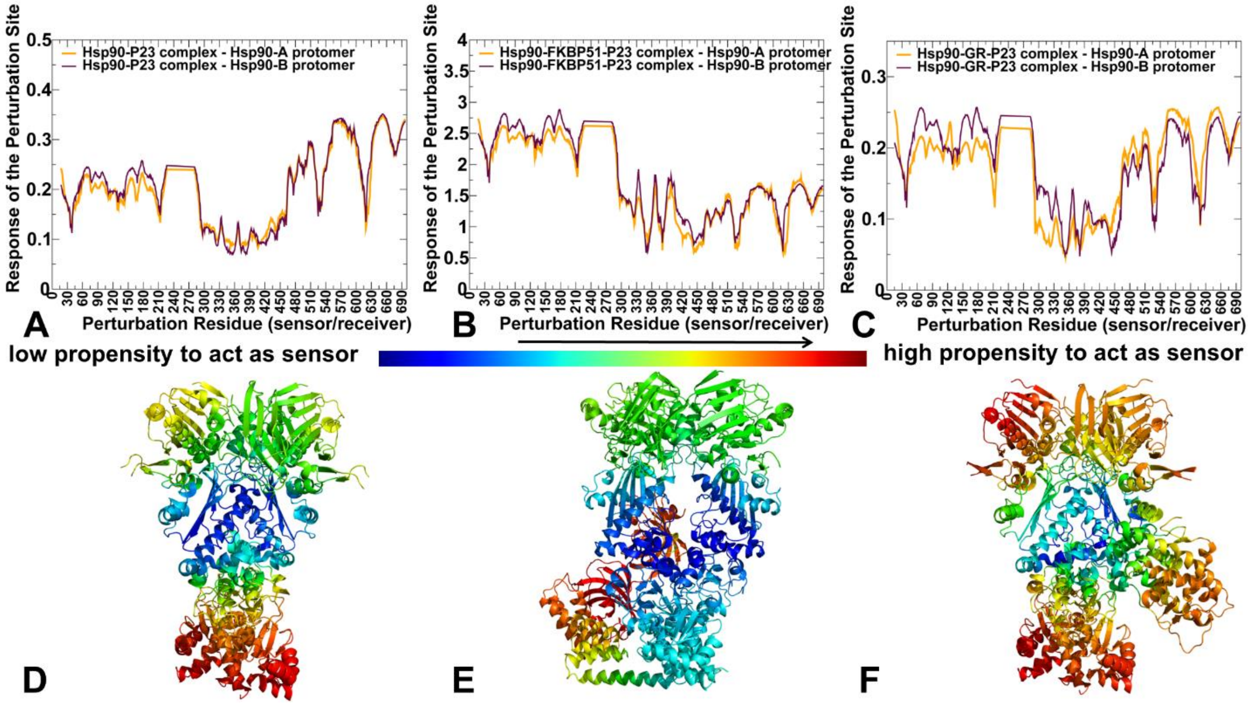


**Figure S4.** PRS analysis and the sensor/receiver residue profiles of the Hsp90 regulatory complexes. The residue-based PRS sensor profiles of the Hsp90 protomers in the Hsp90-P23 complex (A), Hsp90-FKBP51-P23 complex (B), and Hsp90-GR-P23 complex (C). Hsp90-A protomer is in orange lines, Hsp90-B protomer is in maroon-colored lines. Structural maps of the PRS sensor profiles for the Hsp90-P23 complex (D), Hsp90-FKBP51-P23 complex (E), and Hsp90-GR-P23 complex (F). The structure is colored according to the sensor/receiver residue potential with blue-to-red color spectrum corresponding to the increase in the sensor propensities.

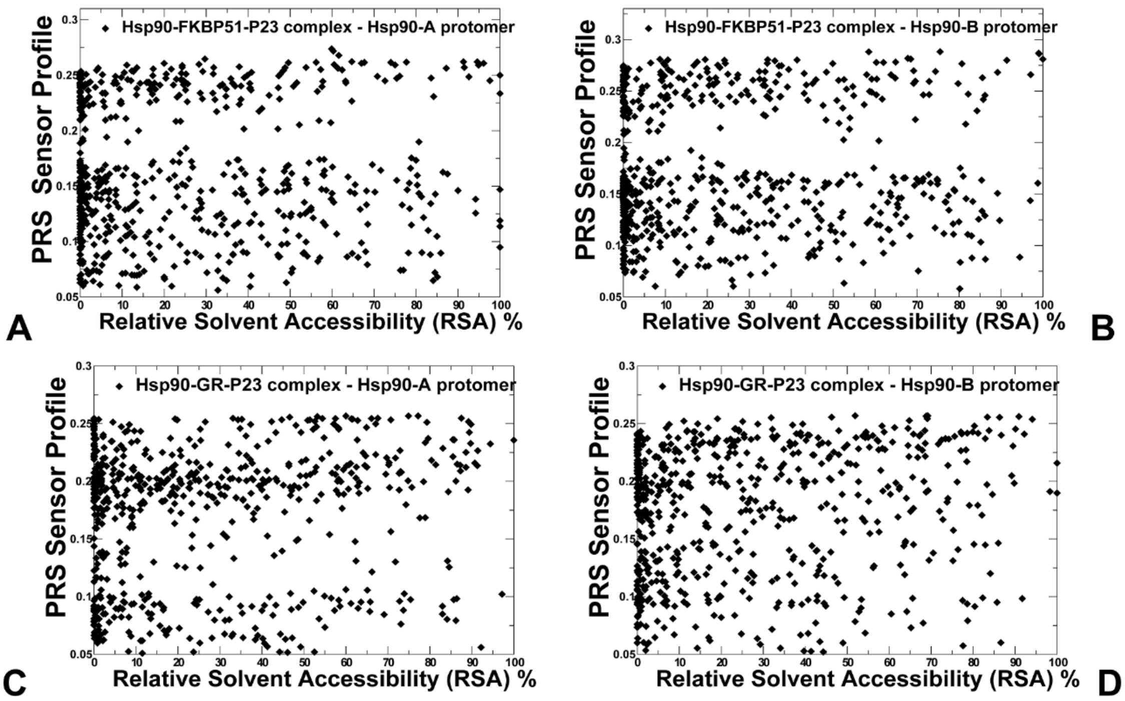


**Figure S5.** The scatter graphs between the PRS sensor profile values and the RSA parameters for the Hsp90 protomer residues in the Hsp90-FKBP51-P23 complex (A,B) and Hsp90-GR-P23 complex (C,D). The residue-base data points are shown in black-colored filled diamonds.
